# ecPath detects ecDNA in tumors from histopathology images

**DOI:** 10.1101/2024.11.13.623494

**Authors:** Mudra Choudhury, Lihe Liu, Anamika Yadav, Owen Chapman, Zahra Ahmadi, Raneen Younis, Chinmay Sharma, Navansh Goel, Sunita Sridhar, Rishaan Kenkre, Aditi Dutta, Shanqing Wang, Eldad Shulman, Saugato Rahman Dhruba, Danh-Tai Hoang, Kevin Tharp, Megan Paul, Denise Malicki, Kevin Yip, Eytan Ruppin, Lukas Chavez, Sanju Sinha

## Abstract

Circular extrachromosomal DNA (ecDNA) can drive tumor initiation, progression and resistance in some of the most aggressive cancers and is emerging as a promising anti-cancer target. However, detection currently requires costly whole-genome sequencing (WGS) or labor-intensive cytogenetic or FISH imaging, limiting its application in routine clinical diagnosis. To overcome this, we developed ***ecPath*** (<u>ec</u>DNA from histo<u>path</u>ology), a computational method for predicting ecDNA status from routinely available hematoxylin and eosin (H&E) images. ecPath implements a deep-learning method we call transcriptomics-guided learning, which utilizes both transcriptomics and H&E images during the training phase to enable successful ecDNA prediction from H&E images alone, a task not achievable with models trained on H&E images only. It is trained on more than 6,000 tumor whole-slide images from the TCGA cohort with the best performance in predicting ecDNA status in brain and stomach tumors (average AUC=0.78). ecPath revealed that ecDNA-positive tumors are enriched with pleomorphic, larger and high-density nuclei. Testing in an independent cohort, ecPath predicted ecDNA status of 985 pediatric brain tumor patients with an AUC of 0.72. Finally, we applied ecPath to identify ecDNA-positive tumors in the TCGA cohort for which no WGS data were available. Like WGS-based ecDNA-positive labels, the predicted ecDNA-positive status also identify poor prognoses for low grade glioma patients. These results demonstrate that ecPath enables the detection of ecDNA from routinely available H&E imaging alone and help nominate aggressive tumors with ecDNA to study and target it.

## Introduction

Circular extrachromosomal DNA (ecDNA) contributes to tumor heterogeneity, progression^1–4^ and resistance to therapy^5^, making targeting ecDNA biogenesis and function a promising strategy to improve therapeutic outcomes. Recent evidence also points to its role in transformation of pre-cancer to cancer, where ecDNA can develop early in the transition from high-grade dysplasia to cancer, and then progressively form and evolve under positive selection^3,6^. Multiple small molecules are in early clinical trial stages to target ecDNA ^7–9^. Detecting the presence of ecDNA in patient tumors can serve as an informative cancer marker for prognosis, patient stratification and patient recruitment to clinical trials. Recent computational and sequencing advancements have enabled the identification of ecDNA from whole genome sequencing (WGS) ^10^. However, WGS is not routinely performed on patient tumors at time of diagnosis due to its high cost. Consequently, there is a critical need to develop alternative methodologies for the detection of ecDNA in clinical settings.

Advances in computer vision have enabled leveraging rich and routinely obtained biomedical images for diagnostic purposes. Digital Pathology especially advanced over the last decade, where computational models distinguish cancer from normal tissue^11^, identify tumor subtypes^12^, detect metastasis^13^ and predict multiple biomarkers^11^ including microsatellite instability^14^, and predict partial transcriptomic and methylation profiles in tumor samples^15,16^. Foundation models, models pre-trained on thousands of hematoxylin and eosin (H&E) images, such as UNI^12^ and CONCH^17^, have enabled transition from application specific models to general-purpose models applicable to multiple clinical tasks.

Developing an image-based approach to detect ecDNA would provide a scalable and accessible diagnostic solution for clinical settings. Previous work focused on predicting ecDNA in single-nucleus karyotypes using high-resolution microscopy^18^. This approach has limited applicability for routine diagnostics due to requirement of laborious cell assays and microscopy instruments typically not available in clinics. If possible, predicting ecDNA from routine biomedical images that are both scalable and accessible offers a potential solution to this challenge. Complementary to this, a previous study has successfully predicted ecDNA status from tumor transcriptome^19^. Building on these advances in computer vision and the established link between transcriptomics and ecDNA, we sought to detect ecDNA status directly from routine pathology slides by inferring tumor transcriptional patterns.

In this study, we present ***ecPath*** (**ec**DNA from histo**path**ology slides), a computational tool to predict ecDNA status from H&E images. ecPath is trained to first predict the tumor transcriptome from an H&E slide and then to determine ecDNA status from the predicted expression profile. This intermediate expression-prediction step by learning on large-scale matched transcriptomics data enabled successful ecDNA detection, which image-only methods cannot (details in Benchmarking Analysis, **Figure 4f**). We first pre-trained ecPath on 6,262 slides from 6,189 patients across 16 tumor types in The Cancer Genome Atlas (TCGA). For 766 patients (797 slides), we obtained ecDNA labels derived from whole genome sequencing (WGS) data from the pan-cancer analysis of whole genomes (PCAWG) (N = 207 slides are from ecDNA positive tumors)^1,10,20^. We validated ecPath’s performance in cross-validation and tested on an independent cohort of primary pediatric brain tumors (N=985) from the Children’s Brain Tumor Network (CBTN)^21^, and provide a resource of predicted ecDNA status for previously unlabeled TCGA samples. Collectively, ecPath offers a novel approach for detecting ecDNA and high-risk cancer patients from H&E tissue slides without the need of DNA sequencing or other molecular profiling techniques.

## RESULTS

### Overview of ecPath pipeline

ecPath is a computational pipeline designed to identify ecDNA from H&E images (**Figure 1a)**. <u>We hypothesized that morphological features visible in H&E slides can predict tumor transcriptomic patterns that are indicative of ecDNA amplifications. This hypothesis builds on previous work demonstrating that ecDNA status correlates with specific transcriptomic signatures</u>^19^, <u>as well as emerging evidence that tissue morphology captured in H&E slides contains information predictive of gene expression patterns</u>. Accordingly, the input of ecPath is an H&E tissue slide and the output is the probability of the tumor sample to be ecDNA positive. ecPath comprises four steps (details below***, Methods 1.1-2.3***): (Step 1) Preprocessing, which involves processing a raw H&E image; (Step 2) Feature Extraction, where morphological features are extracted using the pre-trained foundation model UNI; (Step 3) Transcriptome Modelling (thus called, transcriptome-guided learning), which models the tumor’s transcriptomic profile from the extracted features; and, (Step 4) ecDNA Prediction, which predicts ecDNA from the modelled transcriptome.

**Figure 1:**
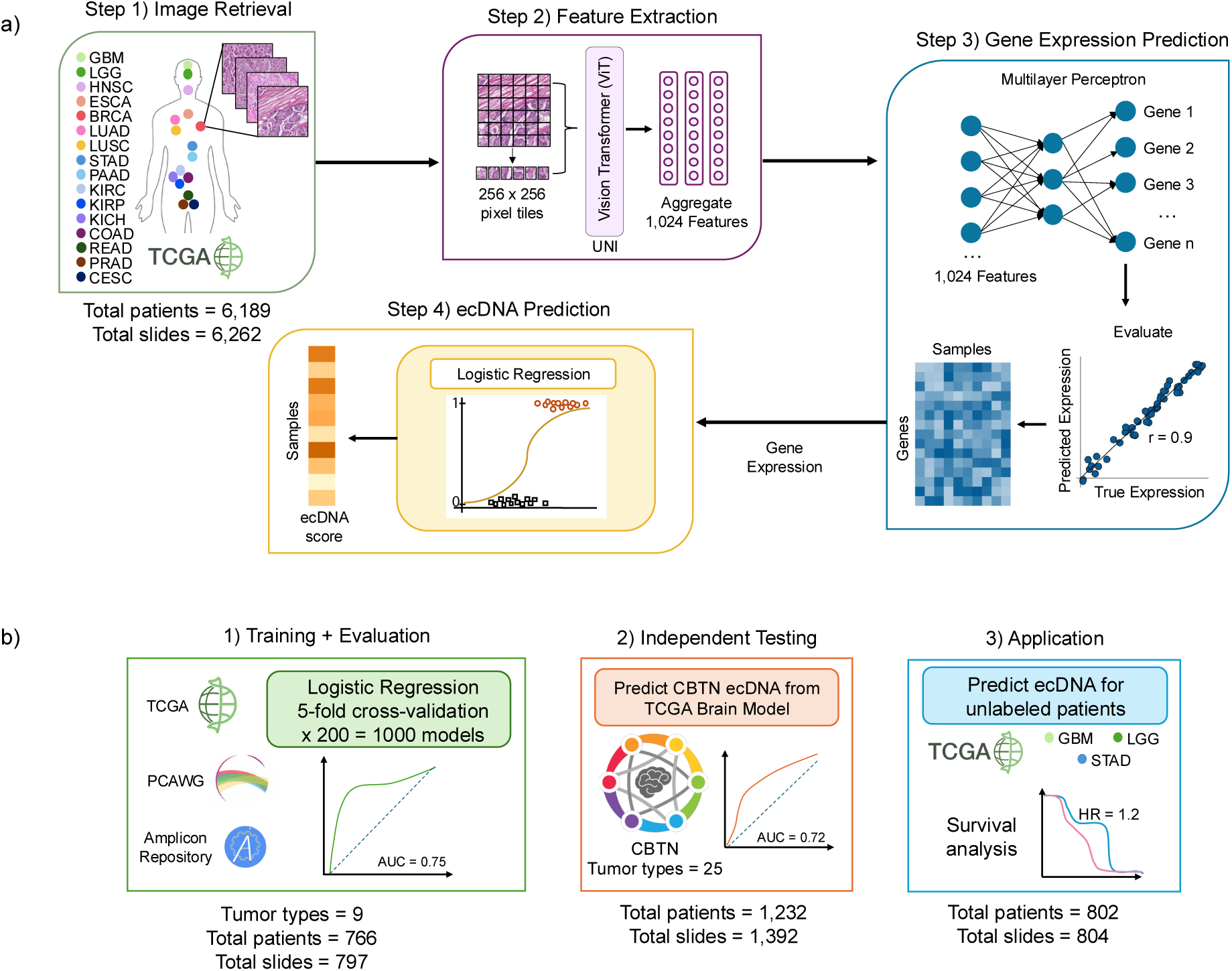
Overview of ecPath and study outline. **a)** Four steps of the ecPath pipeline. Step 1) H&E images are extracted from TCGA from 16 major tumor types and pre-processed. This includes masking, color normalization, and dividing the slide into 256 x 256 images pixel tiles. Step 2) Morphological features are extracted from each tile using a pre-trained foundation model (UNI) yielding 1,024 features for each tile. Step 3) Using these morphological features, ecPath predicts the gene expression profile of individual tumors. This step uses a multi-layer perceptron (MLP) model consisting of three layers: an input layer of features from UNI, a hidden layer, and an output layer with nodes representing the expression for each gene. Genes where true and predicted expression are correlated (Pearson *r* > 0.4) are selected for subsequent ecDNA prediction (Methods 3.1). Step 4) Predicted gene expression is used to build a logistic regression model to determine the ecDNA status of a tumor. ecPath models were trained independently for each cancer type. Pearson correlation is used for correlation analysis and FDR is controlled using the BH method. **b)** ecPath study overview. 1) ecPath models were trained using images and ecDNA annotation from TCGA cohort and PCAWG cohort, where we called ecDNA using WGS. For each tumor-type, a logistic regression model is trained using repeated nested 5-fold cross validation. Each fold is split into training, validation, and test sets. AUC scores for the test-set evaluate model performance. 2) We built a brain-specific model (TCGA low- and high-grade gliomas only) to predict ecDNA labels in the childhood brain tumor network (CBTN) cohort. Model AUC is calculated based on CBTN ecDNA labels for all samples called using WGS (n = 1,232), and primary tumors (n = 985). 3) We applied models trained on the corresponding TCGA cohorts to predict ecDNA in unlabeled samples for three TCGA cancer types (LGG: low-grade glioma, GBM: glioblastoma, and STAD: stomach adenocarcinoma). Cox proportional hazard models were used to test the diaerence in survival between ecDNA positive and negative samples. HNSC: head and neck squamous cell carcinoma, ESCA: esophageal carcinoma, BRCA: breast invasive carcinoma, LUAD: lung adenocarcinoma, LUSC: lung squamous cell carcinoma, PAAD: pancreatic adenocarcinoma, KIRC: kidney renal clear cell carcinoma, KIRP: Kidney renal papillary cell carcinoma, KICH: kidney chromophobe, COAD: Colon adenocarcinoma, READ: Rectum adenocarcinoma, PRAD: prostate adenocarcinoma, CESC: cervical squamous cell carcinoma and endocervical adenocarcinoma.

ecPath was trained on 6,262 H&E whole slide images from 16 tumor types, where both transcriptomics and H&E images are available in The Cancer Genome Atlas (TCGA). **Table S1** provides lists the total number of samples and patients per tumor type. Tumor transcriptomics and ecDNA labels (ecDNA positive vs. negative) were extracted from Xena browser^22^ (transcriptomics) and from Amplicon Repository (ecDNA). We called ecDNA labels using Amplicon Architect to whole-genome sequencing data in PCAWG-cohort.^10,20,23^

In the first step (*Preprocessing*), H&E slides are preprocessed according to the protocols outlined in the current state-of-the-art H&E images to transcriptomics predictor^15^, including background white-space removal, color normalization and partitioning into 256 x 256 pixel tiles. We next obtained morphological features for each tile using UNI, a foundation model based on vision transformer architecture and pre-trained on 100,000 H&E whole slide images (WSI) and >1 million tile images spanning 20 tissue types^12^ (*Feature Extraction*), resulting in 1,024 features per tile (**Figure 1a**, ***Methods 2.1***). Next, we trained a multilayer perceptron model to predict the bulk-gene expression of a tumor for 18,000 genes (*Transcriptome Prediction*). Measured gene expression was utilized as labels during training and only genes with predicted vs. measured expression correlation > 0.4 (predictable genes) were kept for the next step **(Methods 2.2).** Our feature selection process identifies genes where both true and predicted expression levels are strongly associated with ecDNA presence **(Methods 2.3, 3.2).** This concordance-based approach ensures we select reliable gene predictors, reducing the impact of noise due to imperfect expression prediction in selecting features. These criteria ensure a focus on relevant features even in the presence of prediction uncertainties. Finally, we developed an ecDNA prediction model using logistic regression on predicted gene expression. The model is trained and tested in repeated five-fold nested cross-validation. ecPath is trained separately for each cancer-type, considering their morphological uniqueness. In subsequent sections we present ecPath performance in cross-validation across different steps and cancer types, in testing on an independent cohort, and, finally, predict ecDNA in a cohort of tumor samples for which previously no ecDNA information was available (**Figure 1b**).

### ecPath predicts tumor transcriptome from H&E images

For a given tumor H&E slide, ecPath first predicts the transcriptome profile as an intermediate output. Accordingly, we predicted the transcriptome profile of the TCGA cohort, comprising 6,262 H&E slides from 1,689 patients across 16 cancer types (**Figure 2a**, ***Step 3*, *Methods 2.2***)^15^. We consider a gene to be predictable if Pearson’s *r* > 0.4 between predicted and observed expression and FDR-corrected P < 0.05 (concordant with previous methods^15^). Across these 16 cancer types, ecPath successfully predicted the expression of an average of 7,074 genes, with LGG having the highest number of successfully predicted genes (N=12,510). The number of genes for each cancer type is provided in **Figure 2a**. The number of predictive genes in a cancer type and mean performance moderately increases with training set size (*r* = 0.38, **Figure S1a**). We also assessed two H&E image feature extraction methods to evaluate which is superior and provides features that are more predictive of expression: 1) ResNet-50^24^, a convolutional neural network architecture pre-trained on natural images from ImageNet^25^, a method commonly used in digital pathology^15^, vs. 2) UNI, a pre-trained vision transformer pre-trained on 100,000 slide images^12^. We found that UNI-extracted features predict significantly more genes when compared to that from the ResNet-50 method (Mean N = 7,074 vs. 4,084 across 16 cancer types, respectively, **Figure 2b, Figure S1b**). We thus chose UNI for our ecPath architecture, and the transcriptomics predicted using UNI-derived features for subsequent analyses. **Table S2** summarizes the total number of genes regressed, median correlations, and significant genes predicted by ecPATH across TCGA tumors.

**Figure 2:**
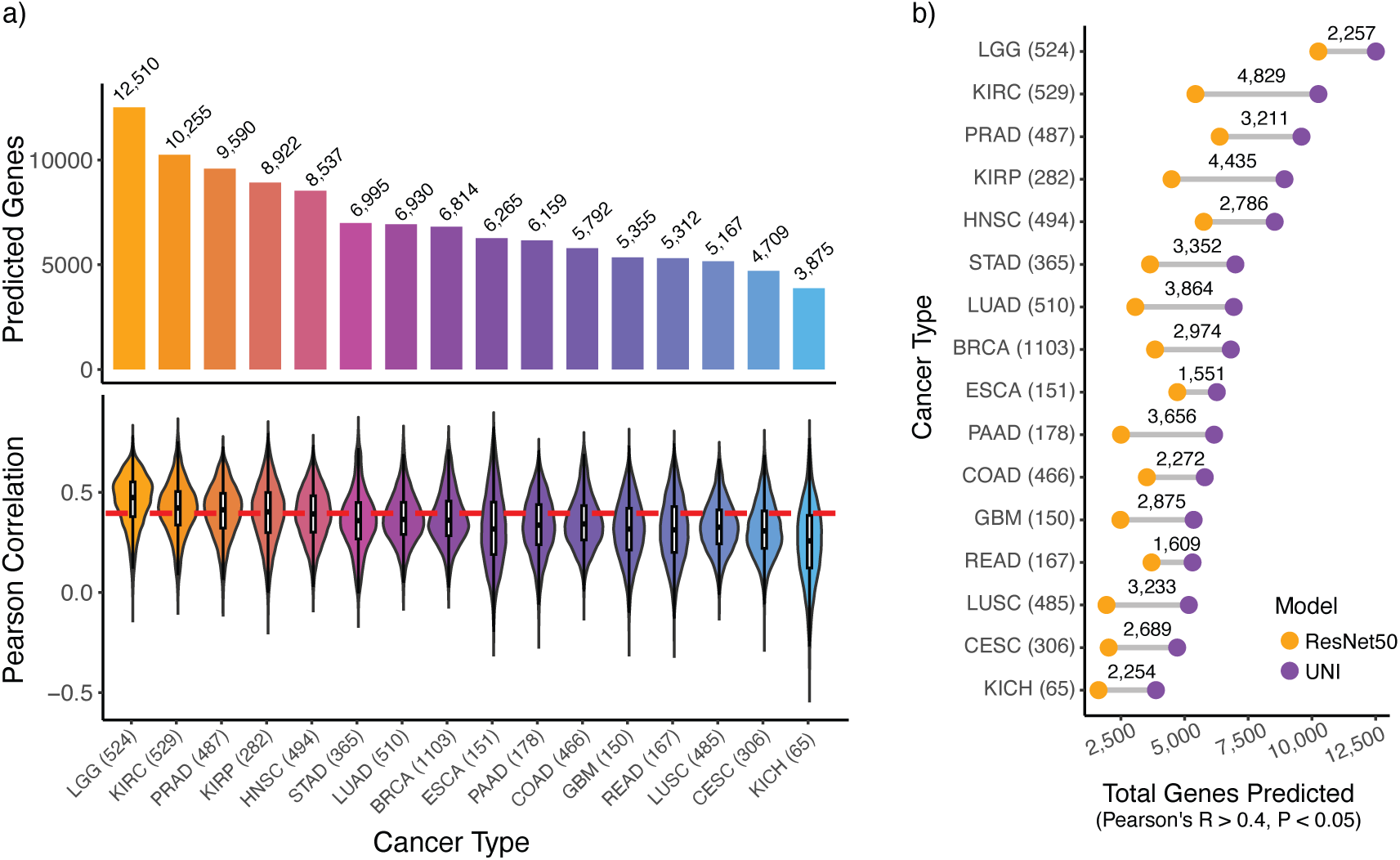
Evaluation of predicted gene expression from ecPath. **a)** The UNI feature extraction protocol and MLP were utilized for gene expression prediction. Total number of significant genes that can be predicted in 16 TCGA tumor types (*r* > 0.4, q < 0.05 with true gene expression) (top). Distribution of Pearson correlation coeaicients across 16 tumor-types (bottom). Red dashed line denotes a correlation threshold of 0.4 **b)** Total number of genes predicted in TCGA through Resnet50 (orange) or UNI (purple) feature extraction methods. Diaerence in total genes predicted per cancer-type are provided above the gray lines.

### Finding cancer types with footprints of ecDNA in transcriptome profile

To train and assess ecPath’s ability to predict ecDNA status, we next obtained ecDNA labels for 797 H&E slides for 766 patients across nine tissues (see **Methods 1.1, 3.2**)^10,23^. Using WGS profiles from Pan-Cancer Analysis of Whole Gnomes (PCAWG)^20^, via Amplicon Architect suite^10^, we called structural variants in this cohort, including ecDNA. The nine tumor types that had at least 10 ecDNA-positive and ecDNA-negative samples were selected for analysis (**Table 1**).

**Table 1:**
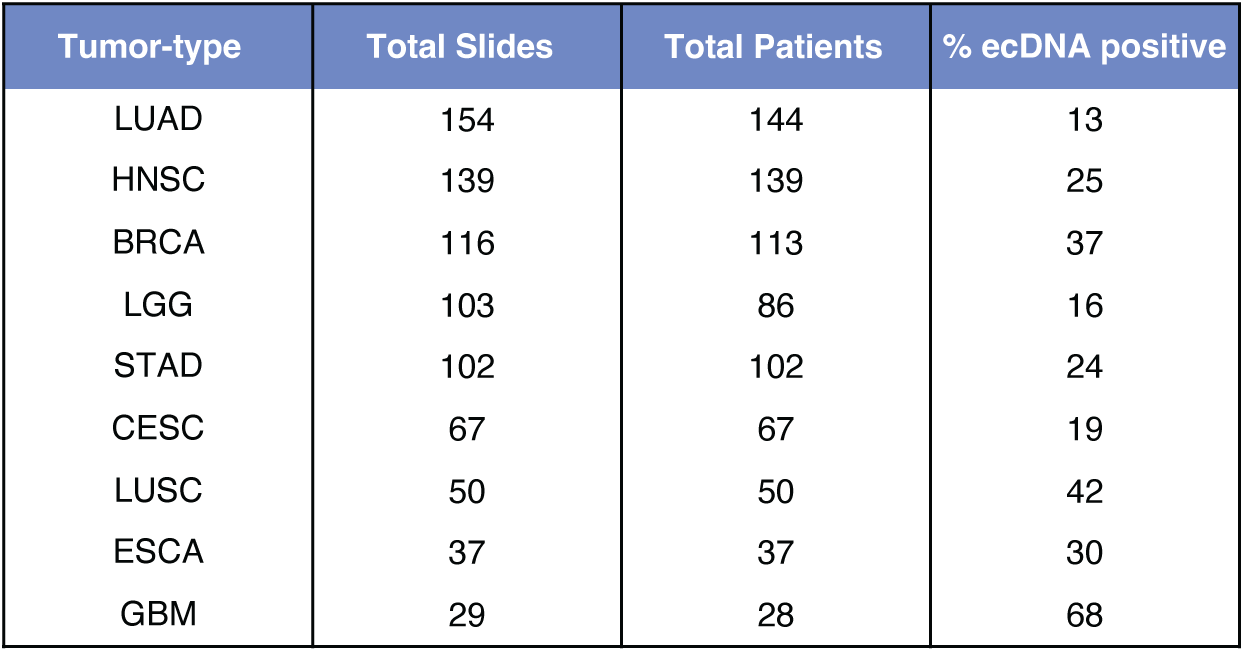
Total number of slides from tumors with known ecDNA status. Total slides, patients, and percent ecDNA positive patients in nine TCGA tumor-types with suaicient slides for prediction.

Following a previous finding that the *measured* tumor transcriptome^19^ can predict ecDNA status^19^, we hypothesized that ecPath’s predicted transcriptome, would be able to predict the same. To this end, we <u>first identified the tumor types where ecDNA footprints can be</u> <u>found in the transcriptomics, by finding cancer types where ecDNA status is predictive</u> <u>using available sample-matched measured transcriptome (RNA-seq)</u>. To this end, for each tumor type, we developed an expression-based ecDNA predictor (**Methods 3.2**). Briefly, classifiers were trained on expression levels of the 150 genes with strongest association with ecDNA (univariate AUC between expression and ecDNA status in training samples). Multiple model types were tested for model-type independent results (logistic regression, SVM, random forest, gradient boosting, and ensemble of the previous four methods). This process was done in five-fold cross-validation, partitioning each cohort into training, validation, and test cohorts, iterated 200 times, resulting in a total of 1000 training cycles **(Methods 3.2).** This analysis revealed that among nine cancer types tested (**Table 1**), the transcriptome profiles of low-grade glioma (LGG), glioblastoma (GBM), and stomach adenocarcinoma (STAD) are predictive of ecDNA status with ROC-AUC of 0.92, 0.83, and 0.81, respectively (AUC > 0.7 as a threshold, **Figure 3a**). Furthermore, this model distinguished ecDNA positive vs. samples devoid of any structural variants (ecDNA, BFB, complex non-cyclic, and linear amplifications, called “no amplicon” here onwards) with an AUC of 0.93, 0.86, and 0.83 in LGG, GBM, and STAD, respectively (**Figure 3b**). We also note that the simplest model, a logistic regression (LR) model, along with gradient boosting (GB) and ensemble models, consistently achieve the highest AUCs (**Figure 3c, Figure S2**). Considering variance-bias tradeoff, we selected the simplest model amongst them – Logistic regression, which was used for all further analysis. This performance is based on measured gene expression and serves as a reference and ceiling (i.e. best possible performance in this cohort) for the performance that can be achieved using predicted expression.

**Figure 3.**
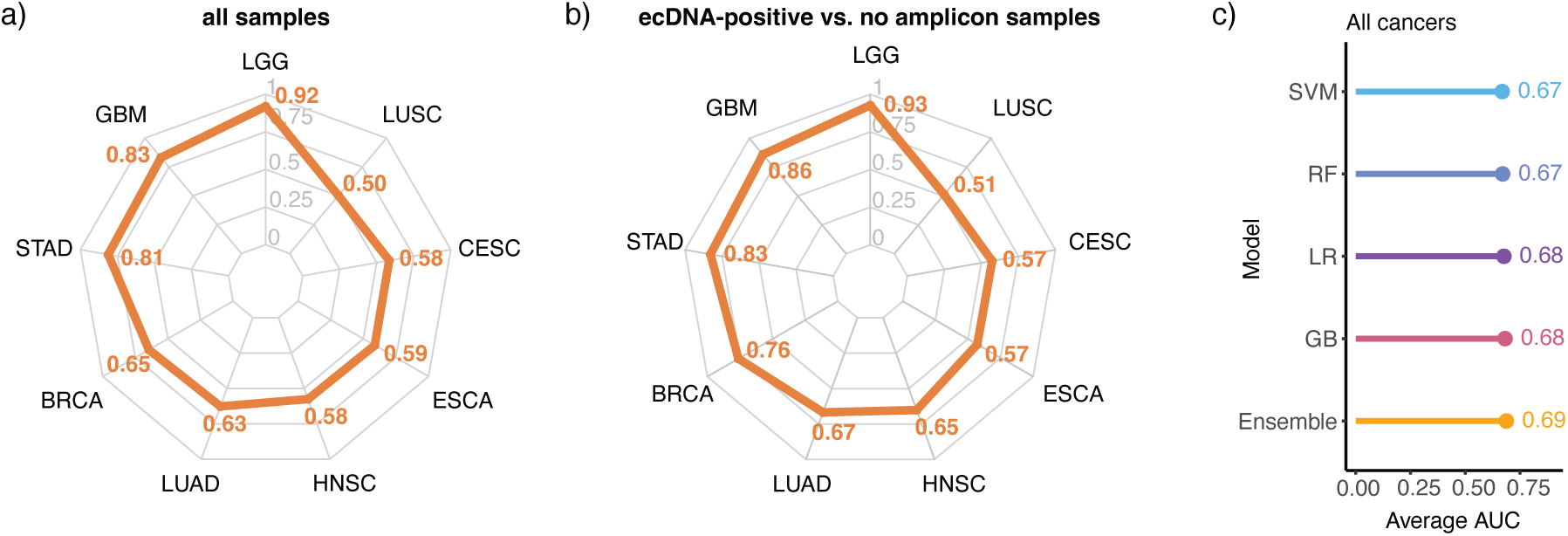
ecPath predicts ecDNA from measured transcriptional profiles. **a)** Area under the curve (AUC) for ecDNA prediction from logistic regression (LR) using true gene expression for nine cancer types. **b)** AUC for ecDNA prediction from true gene expression from nine cancer types for ecDNA positive vs. samples devoid of other amplifications (“no amplicon” samples). **c)** Mean AUC across nine cancer types for each ecDNA prediction model (Methods 3.2). This performance serves as a reference and ceiling (i.e. best possible performance) for the performance that can be achieved using predicted expression. SVM: support vector machine, RF: random forest, LR: logistic regression, GB: gradient boosting, and ensemble of the previous four model.

### ecPath predicts ecDNA status for LGG, GBM, and STAD from the TCGA cohort slides

While the prediction of ecDNA status from known gene expression profiles is valuable, our primary objective was to detect ecDNA solely from H&E images. To this end, we sought to identify a sample’s ecDNA status based on genes whose expression can be reliably predicted from H&E images by ecPath (Pearson’s r > 0.4, q <0.05). Using the same strategy as previously described, we built a logistic regression model using the predicted gene expressions. We again selected genes (N=150) with both measured and predicted expression associated with ecDNA using training samples (***Methods 2.3*, Figure S3**). In cancer types where measured transcriptomics is predictive of ecDNA (i.e. GBM, LGG, and STAD), ecPath can predict ecDNA status, albeit with lower performance (**Figure 4a-c**, AUC = 0.82, 0.78, and 0.73, respectively). ecPath can also distinguish ecDNA vs. other structural variants that are hard to distinguish using sequence-based methods in GBM, LGG and STAD (AUC=0.76, 0.75, and 0.79, respectively. **Figure 4d**). As expected, ecPath’s predicted ecDNA scores for samples that do not contain any types of amplifications (“no amplicon”) were lower than for ecDNA +ve samples and samples that have no ecDNA but other types of amplifications (**Figure 4e**). ecPath performance in all nine tumor-types and the ecDNA prediction scores for each structural variant are provided in **Figure S4a-e**. Performance on nine cancer types was consistent across multiple ML models (**Figure S5a)**.

**Figure 4:**
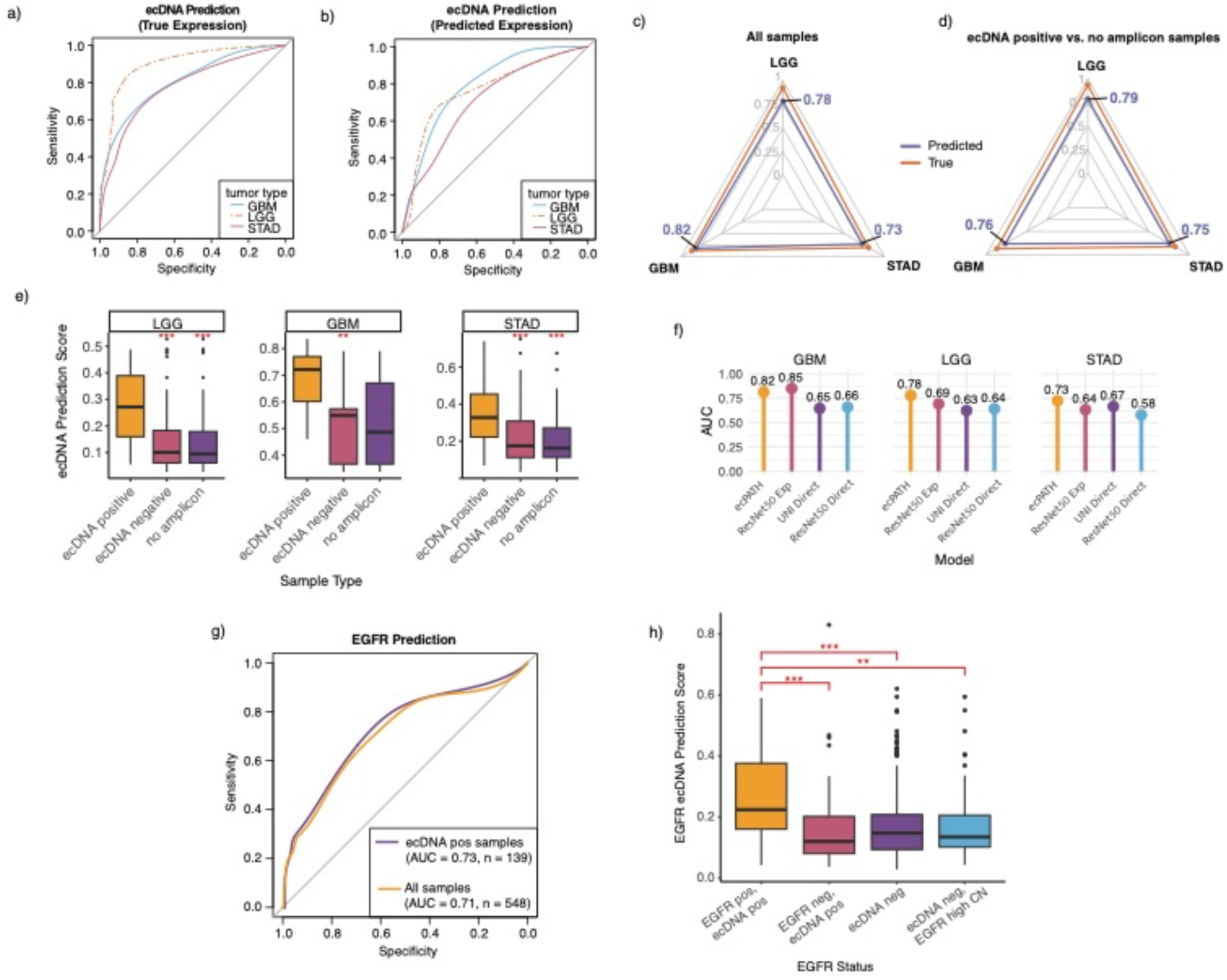
Evaluation of ecDNA prediction. ecDNA prediction ROCs for LGG, GBM, and STAD cancer types trained on true expression **(a)** or trained on predicted expression **(b)**. **c)** Area under the curve (AUC) for each cancer type for models utilizing either true (orange) or predicted expression (purple). **d)** AUC for each cancer type for true (orange) or predicted (purple) expression using only samples that are ecDNA positive and those with no other type of amplifications. **e)** ecDNA prediction probability scores in three TCGA cancer types. Samples are separated into those that are ecDNA positive, ecDNA negative, or a subset of ecDNA negative samples without any other amplifications. Red asterisks denote Wilcoxon rank test significance for the pairwise diaerence in ecDNA prediction score between ecDNA positive samples and either ecDNA negative or no amplicon samples. (*** p < 0.0001, ** p < 0.01, * p < 0.05). **f)** AUC scores of GBM, LGG, and STAD comparing four ecDNA prediction models: 1) ecPath, 2) ResNet50 Exp (utilizing ResNet50 slide features for gene expression prediction), 3) UNI Direct (predict ecDNA status directly from UNI features), and 4) ResNet50 Direct (predict ecDNA status directly from ResNet50 features) (Methods 4). **g)** EGFR ecDNA oncogene prediction ROCs in ecDNA positive samples (purple) or all samples, including ecDNA negative samples (orange). **h)** EGFR prediction probability score for ecDNA positive samples with the EGFR oncogene (EGFR pos, ecDNA pos), ecDNA positive samples without the EGFR oncogene (EGFR neg, ecDNA pos), ecDNA negative samples (ecDNA neg), and ecDNA negative samples that have a high EGFR copy number (ecDNA neg, EGFR high CN). Red asterisks denote Wilcoxon rank test significance as in (e) for the pairwise diaerence between “EGFR pos, ecDNA pos” and the remaining categories.

To validate that our model detects the presence of ecDNA and only the amplification of the harbored oncogenes, we analyzed the ecDNA prediction scores in our cohort, stratified by amplified oncogenes. For the top four oncogenes (EGFR, CDK4, MDM2, and ERBB2) most frequently amplified on ecDNAs in LGG, GBM, and STAD, we found that ecPath detects ecDNA positive from ecDNA negative samples, irrespective of the oncogene present (**Figure S6**) or. vs. ecDNA negative with linear oncogene amplications (Figure 4h). We next tested and confirmed that features used by *ecPath* are directly related to ecDNA biology (details in section “<u>Interpreting ecPath gene expression</u> <u>and morphological features</u>”, **Figure 5A**). This suggests that our ecDNA detection is like a footprint of the ecDNA itself and not just the amplified oncogene.

**Figure 5:**
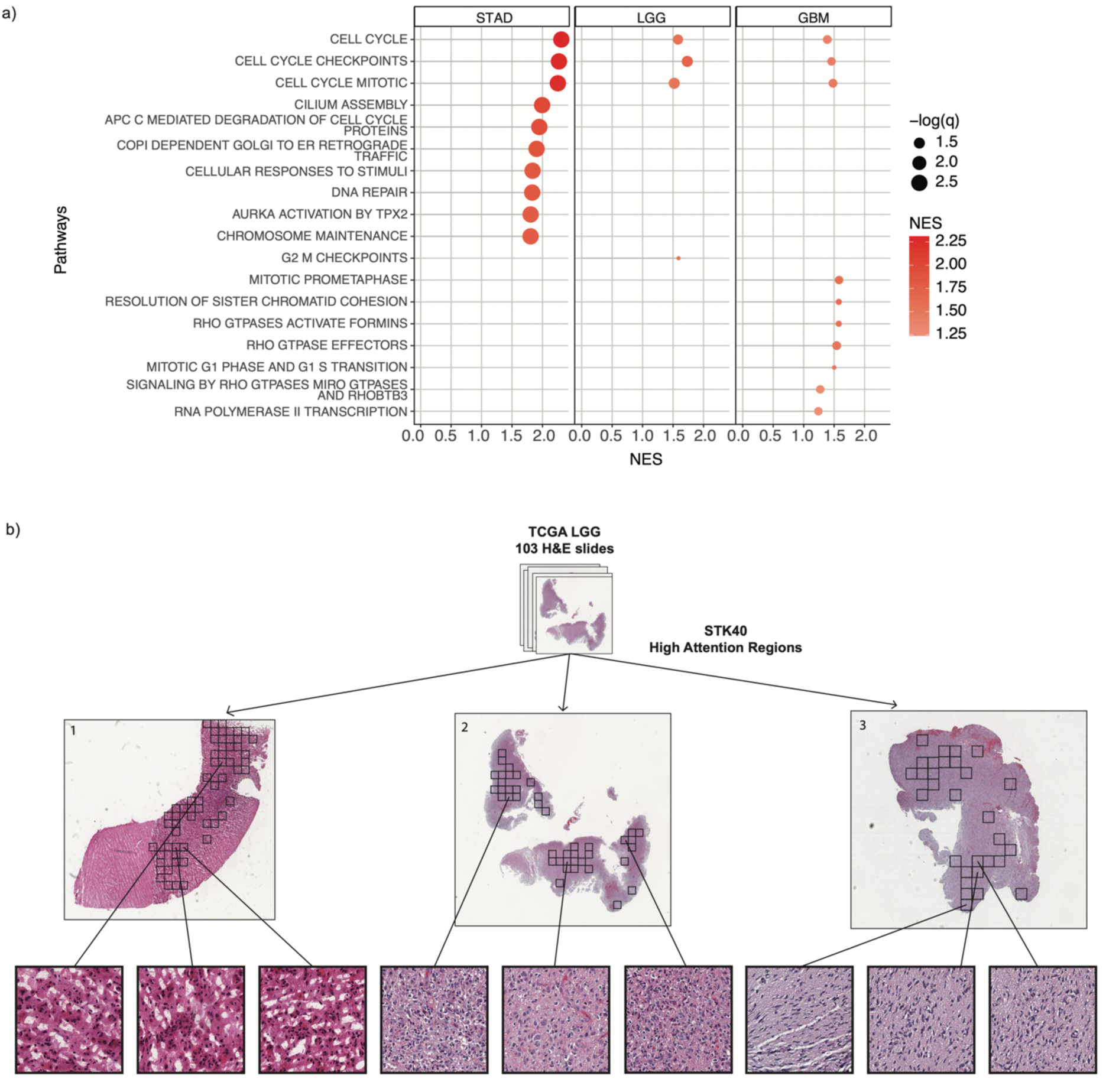
Pathway analysis and Visualization of morphological features. **a)** GSEA of REACTOME pathways in STAD, LGG, and GBM. Size of circles denote the negative log of FDR adjusted p values. Circles are colored by their normalized enrichment score (NES) derived from GSEA. Top 10 pathways are displayed for STAD and GBM. Only pathways with > 20 genes were tested (Methods 6.1). **b)** Visualization of high-attention regions (black squares) for top ecPATH gene STK40 in TCGA LGG slides (Methods 6.2). Slide numbers correspond to pathologist interpretation given in Table S4. Example high-attention tiles are enlarged to demonstrate pleiomorphic and large nuclei and greater density of nuclei, as interpreted by an expert pathologist (Table S4).

#### Benchmarking ecPath

To evaluate ecPath’s performance, we conducted a comprehensive benchmarking analysis. We compared our approach against simpler models that predict ecDNA status directly from slide features, testing both UNI and ResNet50 architectures for feature extraction (**Methods 2.1, 4**). ecPath achieved superior performance with an average AUC of 0.77 across all three cancer types, significantly outperforming the direct prediction models (AUC = 0.63 with ResNet50, AUC = 0.65 with UNI). Notably, the second-best performing model was another indirect approach using ResNet50 (AUC = 0.73, **Figure 4f**), supporting the value of using transcriptional profiles as an intermediate step. ecPath’s superior performance was further confirmed across multiple metrics, including F1 score, precision, recall, specificity, and area under the precision-recall curve (**Figure S5b**).

#### ecPath predicts the oncogene amplified by the ecDNA

Identifying oncogenes amplified on ecDNA can provide valuable biomarkers for cancer diagnosis and targeted therapies. Therefore, we aimed to develop more specialized models capable of predicting ecDNA-amplifications harboring specific oncogenes. Here, we focused on EGFR due to its frequent amplification on ecDNA. We focused on the six of the nine tumor types in which at least one sample had an EGFR-carrying ecDNA (N = 139 samples with ecDNA amplifications harboring EGFR, **Methods 5.1**). We trained a model to predict EGFR positive vs. negative labels (N=24 vs. 115), among ecDNA comprising samples (**Table S3)**, which reached an AUC of 0.73 (**Figure 4g**). As a control, this model was not able to distinguish EGFR-high vs. low cases in tumors without ecDNA (All Samples, **Figure 4g, h**). Testing model specificity to EGFR, we assessed that this model cannot predict CDK4 and MDM2 – two more oncogenes with N>5 amplification (AUC = 0.33 and AUC = 0.41 in CDK4 and MDM2, respectively) (**Figure S7a, b**, **Methods 5.2**). We further tested the prediction probability scores for ecDNA negative samples that have high EGFR copy number (obtained from the Xena browser^22^). EGFR ecDNA probability scores were significantly higher in samples that were positive for EGFR ecDNA than those that were ecDNA negative with high EGFR copy number (**Figure 4h, Figure S7c**, **Methods 5.3**).

#### Interpreting ecPath’s gene expression and morphological features

To interpret ecPath, we first computed pathways enriched in the gene features used by the ecPath logistic regression step (Step 4) in LGG, GBM, and STAD (**Methods 6.1,** *GSEA*^22^ *using REACTOME database*^26^). Concordant with a previous study^19^, genes whose expression were predictive of ecDNA status were involved in cell cycle regulation, DNA damage repair, chromosome organization, and mitotic mechanisms (**Figure 5a**, FDR P < 0.05). We next interpret the tissue regions important to determine the top gene expression features for ecPath. Based on a pathologist’s review, patches that contributed the most in model decision (**Methods 6.2**) are enriched with higher density of nuclei, larger and more pleomorphic nuclei, and larger cells in five LGG slides (**Figure 5b**). Pathologist interpretations of all slides, including those shown in **Figure 5b**, are provided in**, Table S4**. We also find that ecPath can capture expression variance in different parts of the slides^26^ for a few top feature genes with moderate performance (**Methods 6.3, Figure S8**).

### ecPath predicts ecDNA status in an independent cohort of pediatric brain tumor patients

To evaluate ecPath’s performance in an independent cohort, we analyzed 1,392 slides from 1,232 unique patients, spanning 25 major pediatric brain tumor types made available by the Childhood Brain Tumor Network (CBTN, Table S5)^21^. Here, we called ecDNA annotation using available patient-matched WGS data **(Methods 7.1, Data Availability).** To evaluate ecPath performance on this external cohort, we trained a brain-specific ecPath model using all primary brain tumor samples from TCGA, including LGG and GBM. This model predicts expression for 10,768 genes in the LGG and GBM samples (*r >* 0.4, q < 0.05**, Figure S9a-d**) and achieves an AUC of 0.72 in ecDNA prediction across pediatric brain tumor types at the time of diagnosis (n = 985, **Figure 6a-b, Methods 7.1**). These results show that ecPath detects ecDNA in H&E images from an independent test set with comparable accuracy.

**Figure 6:**
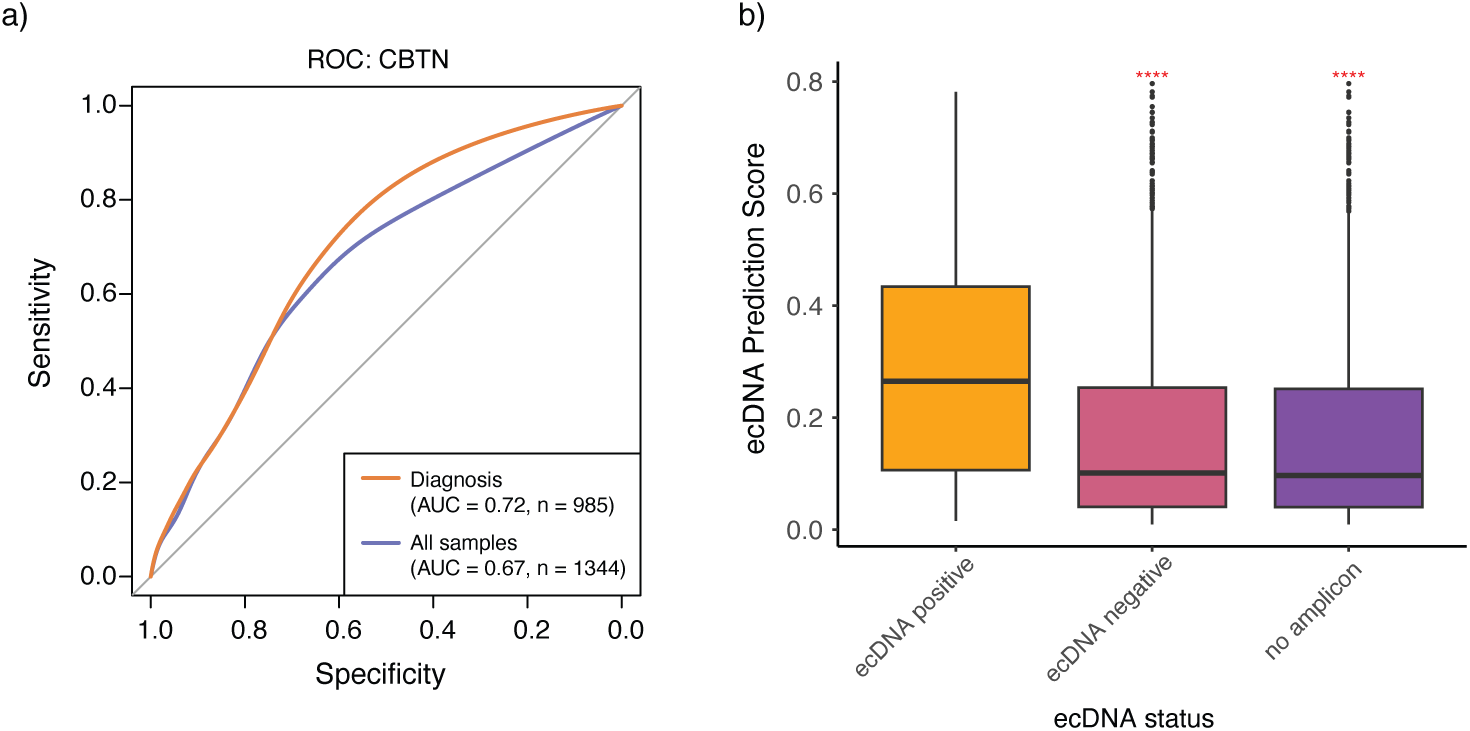
A brain-specific ecPath predicts ecDNA status in independent pediatric brain cohort (CBTN). **a)** ROC of brain LR model of ecDNA prediction applied on CBTN cohort with samples taken at time of diagnosis (orange) or all samples (purple). **b)** ecDNA prediction probability score from brain model for CBTN ecDNA positive (orange), ecDNA negative (magenta), and no-amplicon (dark purple) samples. Red asterisks denote the Wilcoxon rank sum test significance for the diaerence in ecDNA prediction score between ecDNA positive samples pairwise with either ecDNA negative (magenta) or “no amplicon” samples (dark purple). (*** p < 0.0001)

### Predicting ecDNA status for unlabeled TCGA samples

To predict ecDNA status in unlabeled TCGA samples, we applied ecPath to a cohort of LGG (n=421), GBM (n=121), and STAD (n=263) tumors that lack WGS and ecDNA annotations. To this end, we first select a threshold for ecPath’s score to categorize a sample as ecDNA positive that maximizes the sum of sensitivity and specificity (Youden’s J, **Methods 8.1**). Using this threshold, we categorized 802 patient tumors into ecDNA positive and negative samples (positive N=265, negative N=537, **Figure 7a**, **Table S6**). Noting the previous reports that ecDNA is associated with poor prognosis, performed survival regression(s) and found that predicted ecDNA status is significantly associated with poor survival in LGG (**Figure 7b-d**). These results are concordant to a survival analysis of LGG patients in the same cohort (TCGA) where ecDNA labels were determined by conventional WGS data (**Figure 7c,d, Methods 8.2**), suggesting that ecPath can identify high-risk cancer patients marked by ecDNA as a diagnostic and potentially targetable biomarker.

**Figure 7:**
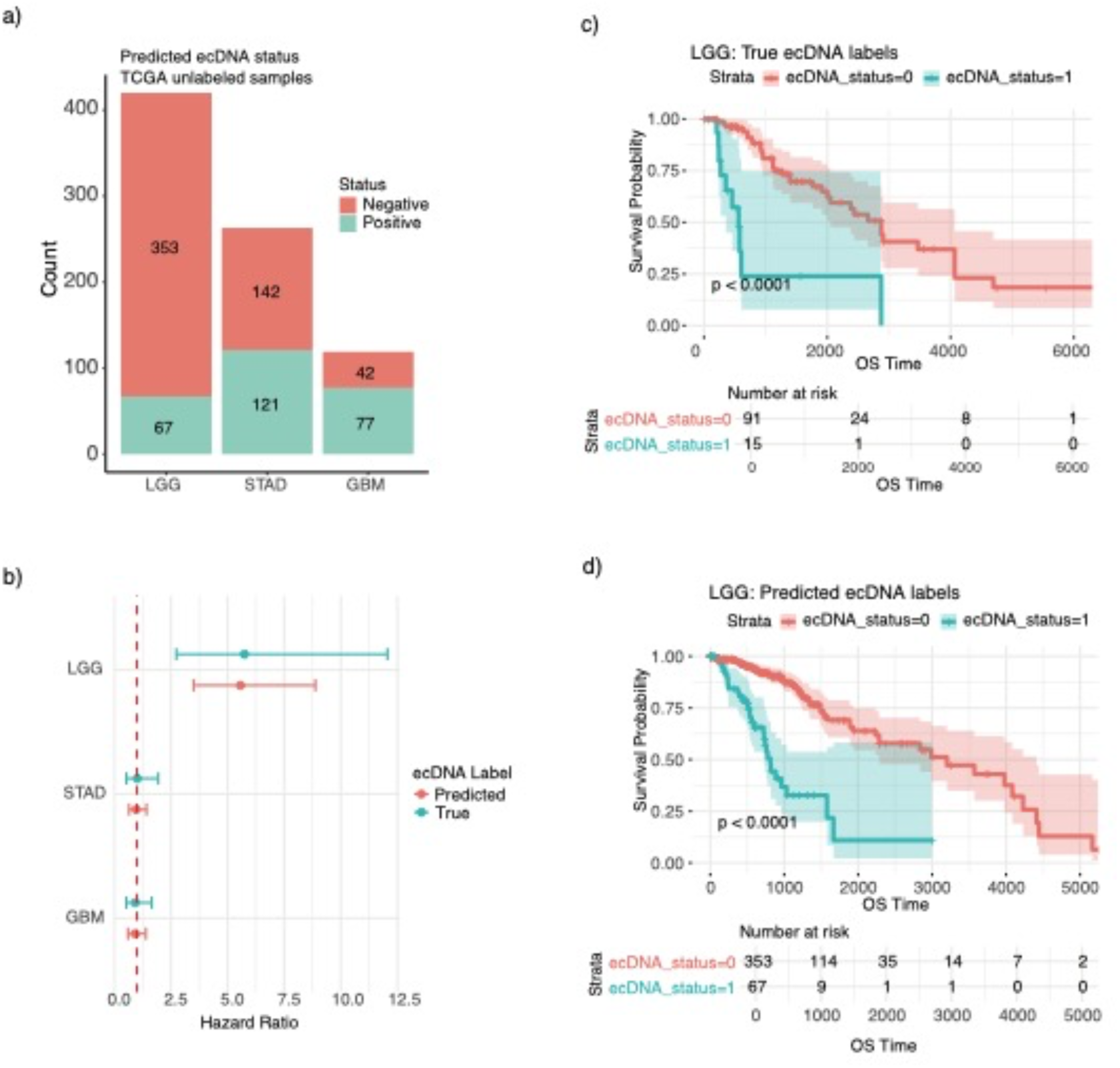
ecDNA status is associated with survival in LGG. **a)** ecDNA predictions for LGG, STAD, and GBM in unlabeled samples. **b)** Multivariate cox-hazard model hazard ratios using true or predicted ecDNA status in three TCGA tumor-types. The error bars represent 95% confidence intervals of the hazard ratio. **c)** Kaplan Meier regression for LGG samples separated by true ecDNA labels. **d)** Kaplan Meier regression for LGG samples separated by predicted ecDNA labels.

## DISCUSSION

We introduce ecPath, a digital pathology method to predict the presence of extrachromosomal DNAs (ecDNAs) from H&E-stained pathology slide images. Two model architecture design choices were instrumental to the performance of our approach: first, the use of a pretrained foundation model to learn biologically relevant features from H&E images; and second, modelling of tumor transcriptome as an intermediate step. To our knowledge, this is the first approach capable of identifying ecDNA status from H&E slides for patients without the need for labor-intensive karyotyping, FISH imaging or DNA sequencing.

We detect footprints of ecDNA in transcriptomes from GBM, LGG, and STAD cancer types, where measured expression is predictive of the ecDNA status. When a brain-specific ecPath model was applied on an independent cohort of pediatric brain tumors, it showed moderate success, demonstrating the model’s broader generalizability to previously unseen examples. When applied to unlabeled TCGA samples, ecPath-predicted ecDNA status associated with poor prognosis for LGG, concordant with WGS-based labels and previous reports^23,27^.

To interpret the features driving ecPath predictions, we visualized key image regions that contributed to gene expression predictions and pathways enriched in ecPath gene expression features (cell cycle and DNA repair categories). The use of the MLP and pre-trained foundation model enabled us to extract gene expression features from the H&E slides. However, more sophisticated models can be developed with larger gold-standard ecDNA datasets, which would enable better model training and validation. Considering our external testing is done in pediatric samples, while yielding comparable performance, underscores the need to test ecPath in larger adult tumor samples to ensure broader applicability. Such data will also facilitate in building models for the prediction of ecDNA copy number or specific oncogene amplifications, which would have significant clinical impact.

Previous research has demonstrated that cancers with elevated ecDNA levels such as GBM and neuroblastoma can exhibit resistance to targeted therapeutics and chemotherapy^29^. Several studies have suggested that the reduction of oncogene-bearing ecDNA molecules may represent a viable therapeutic avenue, as demonstrated through the non-cytotoxic intervention hydrourea in advanced ovarian cancer^30,31^. The ability to predict patient ecDNA status using ecPath can facilitate the identification of ecDNA-positive patients for targeted therapies.

In conclusion, ecPath, provides the first-of-its-kind, scalable and cost-effective solution for the identification of ecDNA-positive patients directly from routine H&E slides. This technique may pave the way for improved understanding of tumor biology, the development of ecDNA-based biomarkers, and the design of more personalized treatment strategies for cancer patients.

## METHODS

### 1. ecPath Training

ecPath development is presented in two parts: Training Data Aggregation and Model Architecture.

#### 1.1. Training Data Curation: Matched Slide, Transcriptomics and ecDNA status from TCGA Cohort

We compiled a total of 6,262 formalin-fixed, paraffin-embedded (FFPE) H&E whole slide images (WSIs) and gene expression profiles from 6,189 patients across 16 tumor-types in TCGA Research Network (https://www.cancer.gov/tcga). All data was extracted from the Genomic Data Commons Data Portal (https://portal.gdc.cancer.gov). Our analysis focused exclusively on primary tumor samples, given the established relevance of ecDNA identification in tumor samples and its previously documented association with patient survival outcomes. To account for potential morphological variations in H&E images across different cancer types, we maintained separate tumor models for each tissue of origin. The complete distribution of slides and patients across tumor types is presented in **Table S1**.

ecDNA labels were available for a subset of the TCGA samples. We obtained ecDNA classifications through the Amplicon Repository PCAWG project^23^, which identified various structural variants (ecDNA, BFB, complex non-cyclic, and linear amplifications) utilizing WGS data from TCGA and the AmpliconArchitect software^10^. A subset of TCGA samples had ecDNA labels across various TCGA tissues (n = 420). Therefore, tumor-types with an insufficient number of samples (< 10 samples in either positive or negative ecDNA labels) were excluded from the analysis. Total patients with ecDNA labels and percent ecDNA positive patients are shown in **Table 1**.

### 2. ecPath Model Architecture

#### 2.1. Feature Extraction via UNI or Resnet

For each H&E image, we implemented slide cleaning and preprocessing procedures as delineated in the DeepPT protocol^15^. In brief, Sobel edge detection^32^ was employed to identify tissue-containing areas, and each 20x magnification WSI was segmented into 256 x 256-pixel tiles. Tiles were excluded if more than 50% of their pixels exhibited low weighted gradient magnitude, as determined by Sobel and DeepPT methodologies^15,32^. RGB color normalization was applied to remaining tiles using Macenko’s method^33^. Following image preprocessing, we utilized the UNI foundation model to extract features^12^. UNI was built via the DINOv2 software, a self-supervised learning algorithm to pretrain vision transformer (ViT) architectures by utilizing student-teacher knowledge distillation^34^. UNI is a large ViT pretrained on the Mass-100K dataset, which consists of millions of tissue patches from 100,426 diagnostic H&E WSIs across 20 major tissue types. We applied the UNI model to extract 1,024 features per 256 x 256 pixel tile across all TCGA WSIs.

For comparative analysis of feature extraction methodologies, we also implemented a second well-established feature extraction methodology. Tiles were resized to 224 x 224 pixels and 2,048 features were extracted from each tile using ResNet50^24^, a convolutional neural network trained on 14 million images from the ImageNet database^25^.

#### 2.2 MLP Regression to Predict Transcriptional Profiles from WSIs

Morphological features extracted from the above step were fed to a multilayer perceptron (MLP) regression algorithm, modeled after the DeepPT^15^ architecture, to predict gene expression. The MLP comprised three layers: (1) an input layer of 1,024 nodes, derived from UNI model features; (2) a hidden layer of 512 nodes; and (3) an output layer with one node per gene. When utilizing ResNet50 for feature extraction and compression, we modified the MLP to have an input layer of 2,048.

The MLP, applied independently to each tumor type, follows the DeepPT framework^15^. We implemented a 5 x 5 nested cross-validation approach, with each outer fold dividing the samples into 80/20 training and testing sets, respectively. To maintain data integrity, images from the same patient were consistently allocated to the same set during cross-validation splits. Five models were trained independently for each internal training and validation set. Final predictions for a gene in the held-out test samples were computed by averaging predictions from the five internal models and across all tiles in the WSI. These final gene expression predictions were utilized to evaluate model performance for each TCGA tumor type and for ecDNA status prediction. The resulting MLP encompassed 25 trained models. To predict gene expression on external cohorts, the average predictions from the 25 pretrained models were used.

#### 2.3. Prediction of ecDNA Status from Predicted Transcriptional Profiles

As a final step of ecPath, we trained a shallow model to predict ecDNA based on predicted expression from our above step. ecDNA status prediction was conducted as described in **Methods 3.2**. Here, top 150 gene features for each model were selected by taking the average univariate AUCs of both true expression and predicted expression in their association with ecDNA status. This was to select for genes that were robustly predictive of ecDNA status in both their true and predicted transcriptional profiles (**Methods 3.2**). Logistic Regression was utilized with the same framework described in **Methods 3.2** to predict ecDNA status from predicted gene expression.

### 3. Evaluating ecPath

#### 3.1 Performance in Gene Expression Prediction

Genes were predicted with sufficient accuracy if they met two criteria: a Pearson correlation coefficient > 0.4 between true and predicted expression, and Benjamini-Hochberg FDR-adjusted p-value < 0.05. These thresholds were established based on the model evaluation component of the DeepPT framework^15^. For subsequent ecDNA prediction, we exclusively utilized the predicted expression of genes that satisfied these significance thresholds.

#### 3.2 Prediction of ecDNA Status using Gene Expression

To evaluate various models for ecDNA prediction based on transcriptional profiles, we built five ecDNA prediction models based on true gene expression: logistic regression, SVM, random forest, gradient boosting, and an ensemble model combining the individual ML methods. Each model was trained separately per tumor type by nested 5-fold cross validation with 200 repetitions, resulting in 1000 trained models for robustness. The samples were split into 80/20 training and testing sets per fold, and training samples were further separated into 80/20 training and validation sets. During cross validation, training and validation sets were utilized for hyperparameter tuning through a grid search methodology for the appropriate hyperparameters. The best model estimators were evaluated by ROC score and utilized for predictions on the test set. To calculate final predictions per test set in the ensemble model, we took a weighted average of the predictions from the four individual ML methods, where the weights were given according to each ML models’ AUC from the validation set. Consequently, the ML models with higher AUCs had greater contribution to final ensemble model predictions.

For each training fold, we utilized rank normalized true expression of 150 genes as features for training. Gene features were selected according to their univariate AUC in predicting ecDNA status, utilizing only samples within the current training fold. The top 150 genes with both the highest and lowest univariate AUCs were considered. We operate under the assumption that genes with high and low AUC values represent those genes that have strongest positive or negative respective association with ecDNA status. We utilized the same framework of ecDNA prediction described above to predict ecDNA status using true expression of only genes that can be predicted with sufficient accuracy by the MLP. In this method, 150 genes are selected from the pool of predictable genes (Pearson’s *r* > 0.4 and adjusted p < 0.05) whose true expression had the strongest association with ecDNA status according to univariate AUC. The true expression of these 150 genes were utilized as features for training. ecDNA prediction based on predicted expression profiles are described in **Methods 2.3**.

### 4. Benchmarking ecPath against other ecDNA prediction methods

We compared ecPath with three other methods for ecDNA prediction. First, we conducted ecDNA prediction as described in **Methods 2.3** replacing gene expression predicted from the ResNet50 feature extractor. We next developed an ecDNA predictor that bypasses the necessity for predicting gene expression from slides, and predicted ecDNA status directly from UNI or Resnet50 features. This model utilizes the MLP algorithm described in **Methods 2.2**. However, the framework was modified to output prediction probabilities for a single ecDNA label as opposed to gene expression. We calculated AUC values in LGG, STAD, and GBM of the ecDNA predictions from the four models: UNI-expression model (ecPath), ResNet50-expression model (ResNet50 expression), UNI direct feature-to-ecDNA prediction model (UNI direct), and ResNet50 direct feature-to-ecDNA prediction model (ResNet50 direct). ecDNA prediction probabilities from all four models were utilized to calculate performance metrics: AUC, F1 score, precision, recall, specificity, and AUPRC. For each tumor-type, F1 scores were calculated by binarizing ecDNA predictions based on an optimal threshold selected through maximizing the sum of sensitivity and specificity (Youden J statistic)^35^.

### 5. Predict EGFR status using ecPath framework

#### 5.1 Training ecPath to predict EGFR in TCGA

We obtained EGFR status from PCAWG^1,20^ in ecDNA positive samples for which we had predicted gene expression. Only tumor-types with at least one EGFR positive ecDNA sample were included, yielding 139 samples (24 and 115 EGFR positive and negative samples, respectively) in six tumor types (LGG, GBM, BRCA, HNSC, ESCA, LUAD, **Table S3**). We used the ecPath LR model framework (**Methods 2.3, 3.2**) to train and evaluate EGFR prediction performance. The model was trained on ecDNA positive samples combined from all six tumor types as there were too few EGFR positive samples in separate tumors for training. Only genes that were predicted in all six tumor types (817 genes) were considered when training. The trained EGFR model was used to predict EGFR prediction probability scores in ecDNA negative samples in the six tumor categories.

#### 5.2 EGFR model performance to predict MDM2 and CDK4 labels

The EGFR model was applied to ecDNA positive samples with known MDM2 or CDK4 labels. Only LGG and GBM tumor types had positive and negative labels for both EGFR and either MDM2 and CDK4. Therefore, the EGFR model was applied to predict MDM2 or CDK4 labels in LGG and GBM tumor types.

#### 5.3 EGFR model performance on ecDNA negative samples with high EGFR copy number

We obtained EGFR copy number from the Xena browser^22^ for ecDNA positive and ecDNA negative samples. Samples were divided into three categories according to EGFR copy number (CN) percentile (low: CN below 25^th^ percentile, medium: CN between 25^th^ and 75^th^ percentile, high: CN above 75^th^ percentile). Wilcoxon rank sum tests were utilized to test the difference between EGFR prediction probabilities between sample categories.

### 6. Interpretation of ecPath model features

#### 6.1 Pathway analysis of gene features for ecDNA prediction

We aggregated gene features selected in the ecPath logistic regression models for LGG, GBM, and STAD. Gene features were ranked according to their average univariate AUC, which was calculated during training of LR models per tumor-type (**Methods 2.3**). We utilized GSEA^36^ to enrich for pathways in the REACTOME database^26^ and restricted pathways to a minimum size of 20 genes.

#### 6.2 Visualize slide-level attention in TCGA LGG

We selected the genes that can be accurately predicted by ecPath, (*r* > 0.7), and are highly predictive of ecDNA status(average univariate AUC > 0.8) as candidate genes for attention analysis. To identify regions of interest that were predictive of candidate gene expression, we built a single-gene MLP prediction model in TCGA LGG, with the implementation of an attention framework based on a previous method of regression-based deep learning^37^. After the identification of high attention regions by our MLP-attention model in the example TCGA LGG slides. We sought the expertise of a pathologist for further interpretation of the regions in both ecDNA positive and ecDNA negative slides identified by our MLP-attention model.

#### 6.3 Visualize spatial gene expression predictions in independent BRCA HER2+ cohort

To evaluate our model’s ability to predict spatial-level gene expression, we applied ecPath to an independent BRCA HER2-positive cohort with known spatial gene expression^38^. This cohort had H&E stained images (n=36) from 8 patients and known gene expression through spatial microarray analysis. We filtered for genes in the BRCA HER2+ cohort that had non-zero variance and more than 1000 microarray spots across all slides. We then selected genes that were used in our BRCA ecDNA prediction model (n = 785 genes) to train single-gene MLP-attention models **(Methods 2.2)** in the TCGA BRCA cohort.

We extracted 512 x 512 pixel regions around each microarray spot in the BRCA HER2-positive samples. We utilized the above trained single-gene models with the attention component to obtain tile-level attention scores and predicted gene expression for 785 genes of interest. For visualization purposes, we showed a top example patient slide with the a candidate gene (SETD6) with the highest concordance between true gene expression and predicted gene expression (Pearson R = 0.518, P =1.8e-08) and an example gene (S100A11) in the same patient with the highest concordance between true expression and attention score for gene expression prediction (**Figure S8**, Pearson R = 0.428, P = 6.5e-07).

### 7. Evaluate ecPath on independent cohort

#### 7.1 ecDNA prediction validation in CBTN

ecDNA labels from WGS and transcriptional profiles were obtained from the CBTN cohort^21^ Kids First portal (Transcriptional profiles: https://kidsfirstdrc.org/, WGS: https://commonfund.nih.gov/KidsFirst). We trained ecPath on both LGG and GBM samples to develop a brain-specific model for the prediction of gene expression and ecDNA status. The MLP model was trained as described as in **Methods 2.2** with the addition of stratifying LGG and GBM samples across folds. We obtained 1,392 H&E slides from 1,232 patients that were part of the CBTN cohort across 25 cancer types (**Table S5**).

A subset of patients had multiple slides, as they were sampled at different disease time-points (diagnosis, progression, and recurrence). In these cases, predictions were done separately per slide. We used the slide preprocessing and UNI feature extraction methodology that was consistent with **Methods 2.1**. UNI derived features and brain-specific MLP model weights were utilized to predict gene expression in the pediatric tumor samples part of CBTN. As the MLP trains 25 models during cross-validation, gene expression predictions per slide were averaged across all 25 pre-trained brain MLP models. We used Pearson’s correlation coefficient and Benjamini-Hochberg corrected p-values to evaluate gene expression predictions in CBTN for all predicted genes that also had matching true expression.

The LR ecDNA prediction algorithm was trained on both TCGA LGG and GBM samples for the purpose of developing a brain-specific ecDNA prediction model. We utilized the ecPath training procedure outlined in **Methods 2.3**. The LGG and GBM combined model achieved an AUC = 0.75 in TCGA brain samples **(Figure S9b).** Pre-trained LR model parameters and gene features for prediction of ecDNA status were applied to predict ecDNA status in the CBTN cohort.

Gene expression was evaluated as in **Methods 3.1**. As expected, the correlation between true and predicted expression was much higher in the TCGA brain samples than in the CBTN sample predictions **(Figure S9ac-d,).** However, a total of 1,032 of the 10,768 genes successfully predicted in the original LGG-GBM brain model were significantly correlated between true and predicted expression in CBTN **(Figure S9d).** Finally, the brain specific ecDNA prediction model, which utilized 988 gene features in the 1000 LR models, was applied to the CBTN cohort to predict ecDNA status.

### 8. Application of ecPath on unlabeled samples

#### 8.1 Predict ecDNA status of unlabeled TCGA samples

We used pre-trained ecPath models for GBM, LGG, and STAD to predict transcriptional profiles and ecDNA status of samples with unknown ecDNA profiles. For each tumor type, the MLP model weights derived from nested 5-fold cross-validation procedure were applied to predict gene expression. Gene expression predictions were averaged across the resulting 25 pre-trained MLP models to obtain final gene expression in the samples with unknown ecDNA labels.

Next, we leveraged tumor-specific LR model parameters and gene features from the 1000 pre-trained LR models to predict ecDNA status. The final prediction scores were the average from 1000 pre-trained tumor-specific LR models. In cases where a patient had multiple slides, prediction scores were averaged across all slides.

For each tumor-type we selected a prediction probability score threshold to binarize ecDNA labels. The selected probability threshold was the value that maximized the Youden’s J statistic^35^, which aims to maximize the difference between the true positive rate (sensitivity) and the false positive rate (1-specificity).

#### 8.2 Patient Survival of TCGA samples

Patient survival data, including overall survival status and survival time, were obtained for the TCGA samples from the Xena browser, NCI Genomics Data Commons^22^. We performed Kaplan-Meier regression analysis and log-rank tests to assess the differences in survival between ecDNA-positive and ecDNA-negative samples. Samples with known and unknown ecDNA status were considered separately. Additionally, we employed the Cox proportional hazards model to calculate the hazard ratios and 95% confidence intervals for the ecDNA-positive samples, using either the true ecDNA labels or the newly predicted ecDNA labels.

## ACKNOWLEDGEMENTS

We thank Sanford Burnham Prebys Medical Discovery Institute for providing start-up funding to the Sinha Lab to support this study and infrastructural support. We thank the Hearst foundation for providing financial support to acquire computational resources required for the study. This project is supported by the UCSD/Rady Children’s Hospital pediatric hematology oncology fellowship program (S.S.), a generous endowment by the Clayes Foundation to the Research Center for Neuro-Oncology and Genomics within the Rady Children’s Institute for Genomic Medicine, a Hannah’s Heroes St. Baldrick’s Scholar Award (L.C.), the Dragon Master Foundation (L.C.), funding from the National Institutes of Health (NIH) National Institute of Neurological Disorders and Stroke Institute R01 NS132780 (L.C.), R21 NS130137 (L.C.) and R21 NS120075 (L.C.), and the NIH National Cancer Institute P30 CA030199-42S1 (L.C. and K.Y.), and F31 CA271777 (O.S.C.). This research was conducted using data made available by The Children’s Brain Tumor Network (formerly the Children’s Brain Tumor Tissue Consortium) and the St. Jude Cloud. We thank J. H. Zhang, C. McLeod, M. Brown and A. Resnick for facilitating data access.

## DATA AVAILABILITY

The H&E images used in the study were downloaded from TCIA webpage for TCGA (https://www.cancerimagingarchive.net/), requested and received access of restricted CBTN cohort via CBTN Request Submission Form (https://portal.kidsfirstdrc.org/). The data generated in this project to run ecPath for independent samples or re-train ecPath is provided at: https://doi.org/10.5281/zenodo.14057816

## CODE AVAILABILITY

The ecPath software package is freely available on Github for academic use, providing three main functionalities: (1) a reviewer mode for replicating model outputs, (2) scripts for reproducing all figures, and (3) a production mode for developing new models using the ecPath architecture. The complete source code and documentation can be accessed at: https://github.com/Sinha-CompBio-Lab/ecPATH

## Contribution

S.S., and L.C. conceptualized the idea of predicting ecDNA from H&E. M.C., S.S., L.C. developed the framework for predicting ecDNA from H&E slides. M.C. led the computational analysis and wrote the first version of ecPath scripts focused on one cancer type. L.L and M.C later expanding the model to multiple cancer types and developing the methodology for identifying important decision-making regions. A.Y. performed independent model validation. while O.C. and C.S. generated ecDNA labels for training and independent testing. O.C., S.Sr., R.K., A.D., S.W., C.S. and M.P. analyzed the CBTN cohort and generated ecDNA labels for training and independent testing. D.M. performed pathological review of selected H&E slides. E.S., S.R.D., D.H., and E.R. provided guidance on the expression prediction analysis. Additional manuscript revisions and feedback were provided by A.Y., Z.A., R.Y., N.G., K.Y., E.S., S.R.D., D.H., and E.R. M.C. and S.S. wrote the initial manuscript, with L.L. and L.C. providing critical revisions.

---

**Figure S1:**
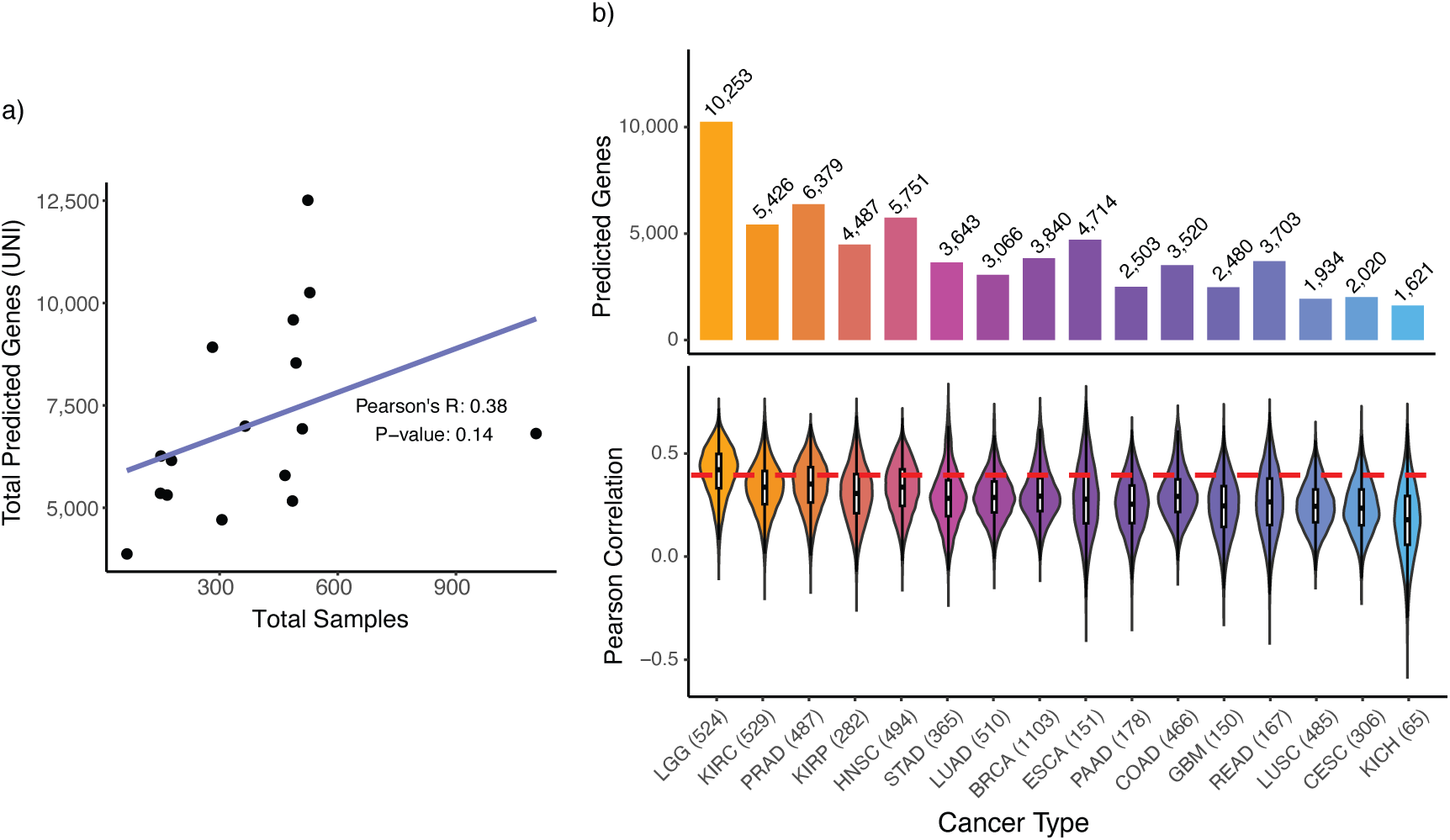
Evaluation of predicted gene expression. **a)** Number of samples in tumor type vs. total genes predicted by UNI for each tumor type. Pearson’s *r* and p-value denotes correlation strength. **b)** Gene expression prediction from ResNet50 and MLP model. Total number of significant genes that can be predicted in 16 TCGA tumor types (Pearson’s correlation R > 0.4, q < 0.05 with true gene expression) (top). Distribution of Pearson correlation coeaicients across 16 tumor-types (bottom).

**Figure S2:**
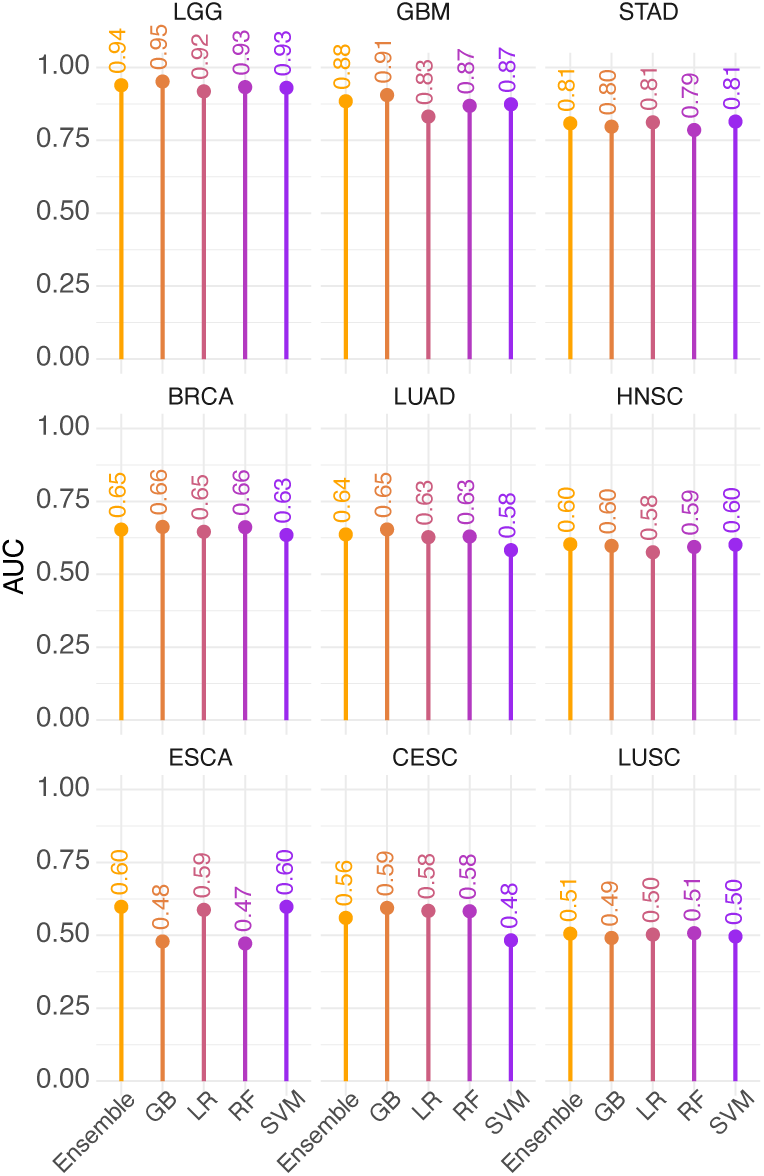
ecDNA prediction performance based on measured expression in four models. Area under the curve (AUC) for four models to predict ecDNA status from measured gene expression profiles for nine cancer-types (Methods 3.2). GB: gradient boosting, LR: logistic regression, RF: random forest, SVM: support vector machine.

**Figure S3:**
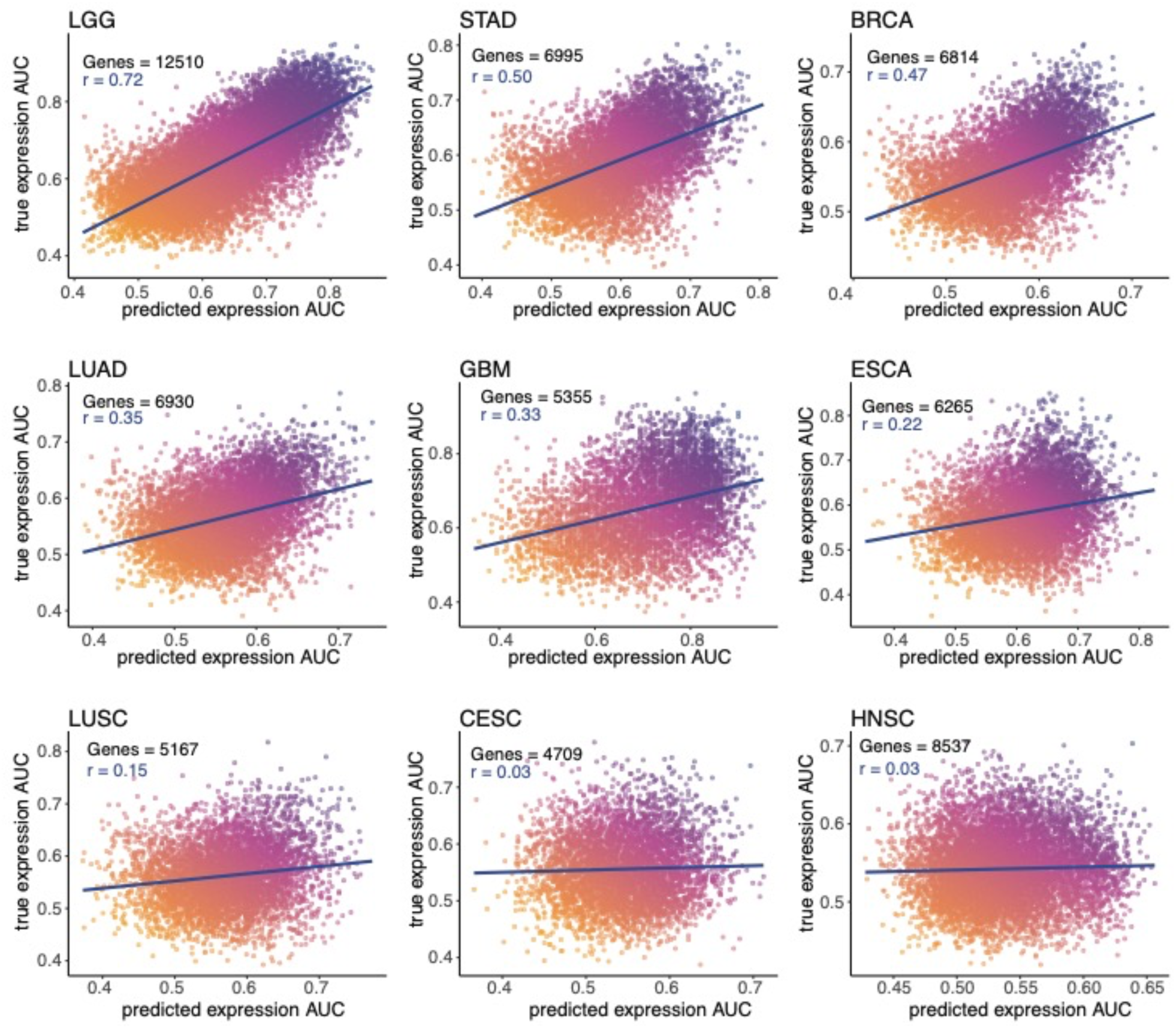
Prediction power of true or predicted transcriptional profiles for ecDNA status. Univariate AUC of true gene expression (y-axis) or predicted gene expression (x-axis) for predicted genes in the MLP model. Pearson R values denote the correlation between AUCs based on predicted or true expression for each gene.

**Figure S4:**
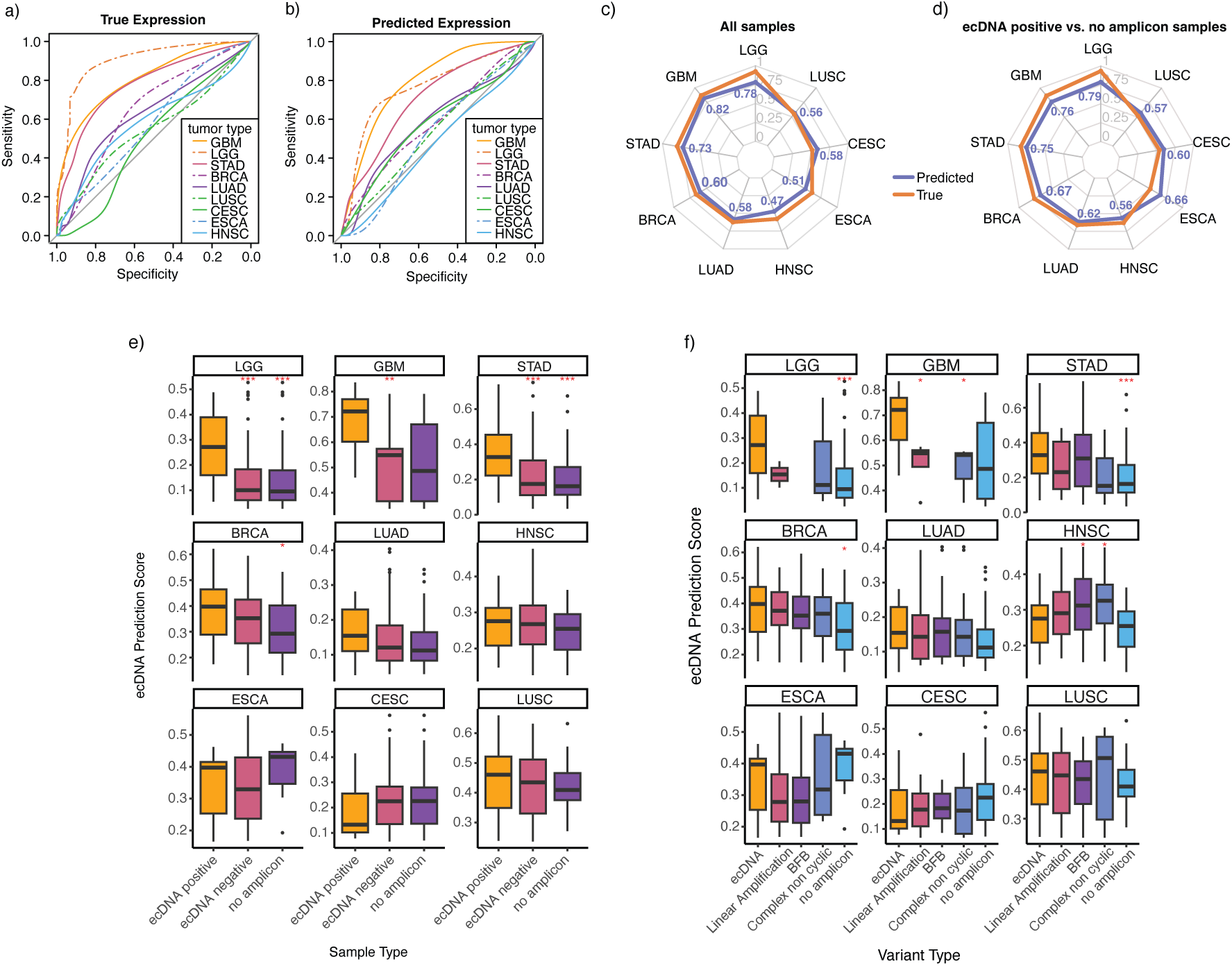
Evaluation of ecDNA prediction in nine tumor types. ecDNA prediction ROCs for nine TCGA cancer types trained on true expression **(a)** or trained on predicted expression **(b). c)** Area under the curve (AUC) for each cancer type for models utilizing either true (orange) or predicted expression (purple). **d)** AUC for each cancer type for true (orange) or predicted (purple) expression using only samples that are ecDNA positive and those with no other type of amplifications. **e)** ecDNA prediction probability scores across nine TCGA cancer types. Samples are separated into those that are ecDNA positive, ecDNA negative, or a subset of ecDNA negative samples without any other amplifications (no amplicon). **f)** ecDNA prediction score of samples categorized as ecDNA positive (ecDNA) or ecDNA negative samples with linear amplifications, breakage fusion bridge (BFB), complex non-cyclic variants, or samples with no other amplifications. Red asterisks denote Wilcoxon rank test significance for the diTerence in ecDNA prediction score between ecDNA positive samples and either ecDNA negative (yellow) or no amplicon (purple) samples. (*** p < 0.0001, ** p < 0.01, * p < 0.05).

**Figure S5:**
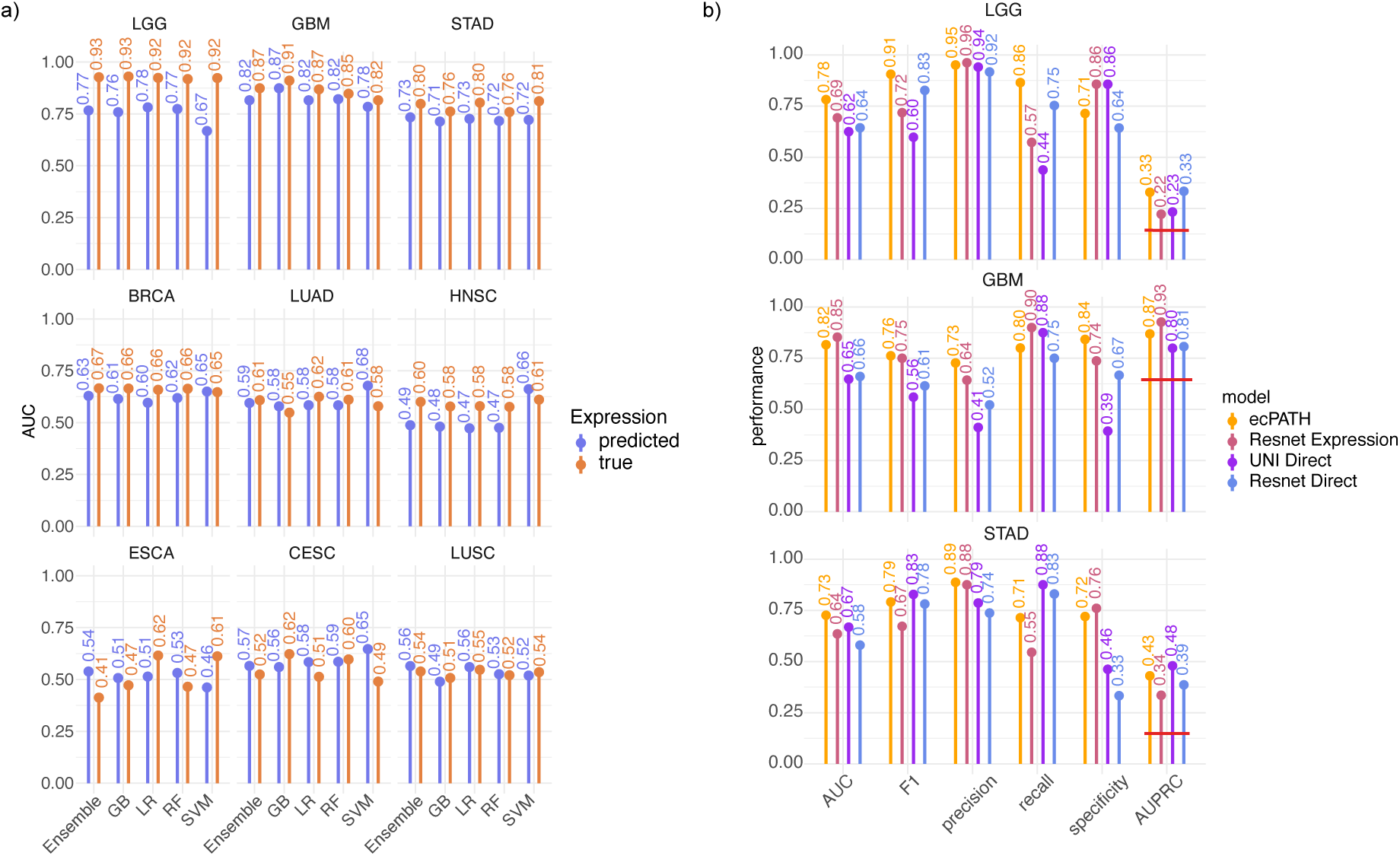
ecDNA prediction performance. **a)** Area under the curve (AUC) for five models to predict ecDNA status from measured gene expression profiles (purple) or predicted expression (orange) for nine cancer-types. Models are an aggregate of the 1000 training cycles, with the top 150 genes selected according to univariate AUCs across training samples (Methods 3.2). GB: gradient boosting, LR: logistic regression, RF: random forest, SVM: support vector machine. **b)** Performance metrics for ecPATH, Resnet Expression, UNI Direct, and Resnet Direct prediction models. Red line on area under precision recall curve (AUPRC) denotes the randomized baseline AUPRC for the tumor type.

**Figure S6.**
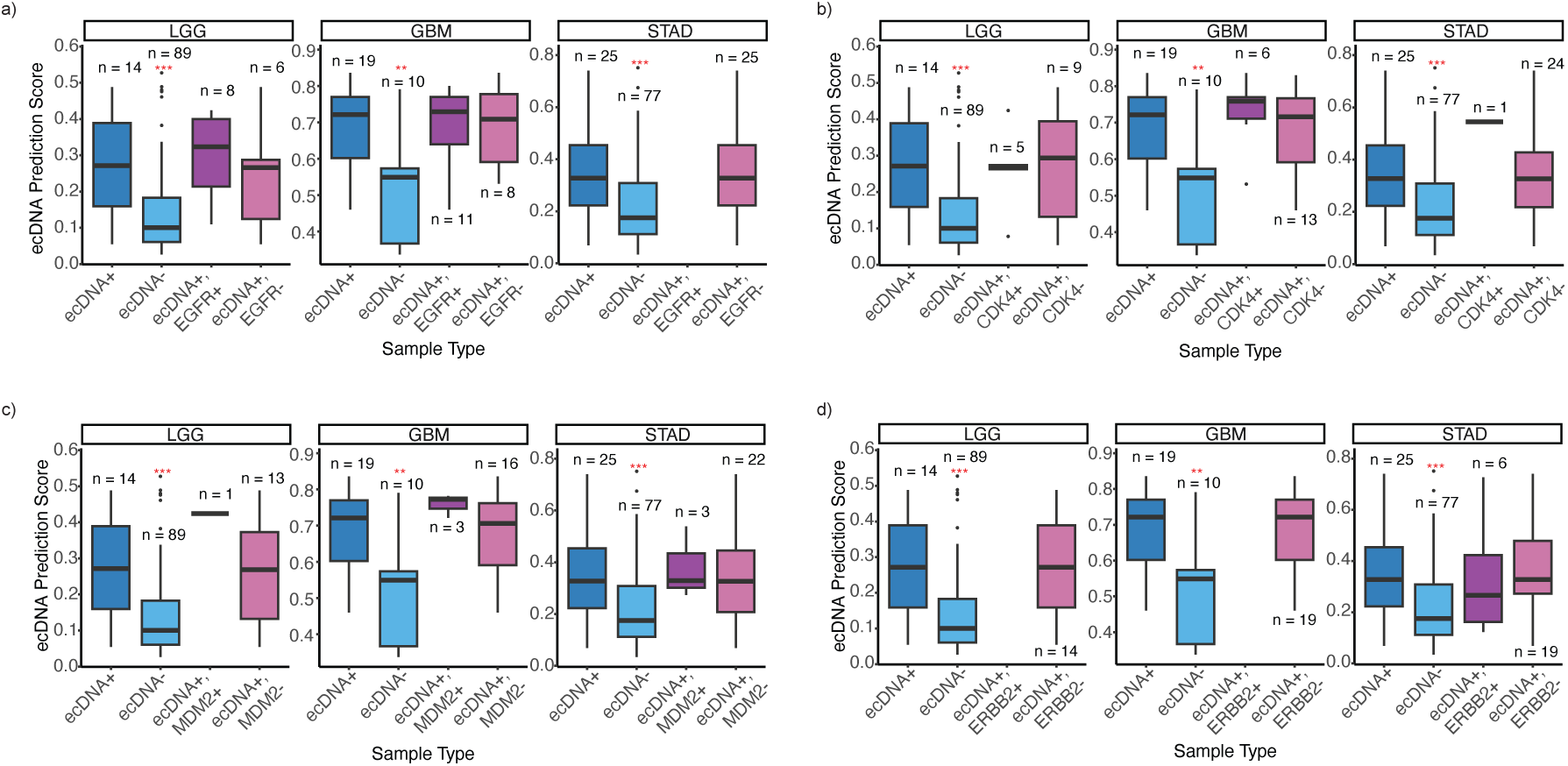
ecDNA prediction scores for samples stratified by oncogene presence. ecDNA prediction score for samples that are ecDNA positive (ecDNA +), ecDNA negative (−), and ecDNA positive samples stratified by presence or absence of the EGFR oncogene **(a)**, CDK4 oncogene **(b)**, MDM2 oncogene **(c)**, ERBB2 oncogene **(d)**. Red asterisks denote Wilcoxon rank test p-value diTerence between sample category and ecDNA positive samples scores (*** p < 0.0001, ** p < 0.01, * p < 0.05).

**Figure S7.**
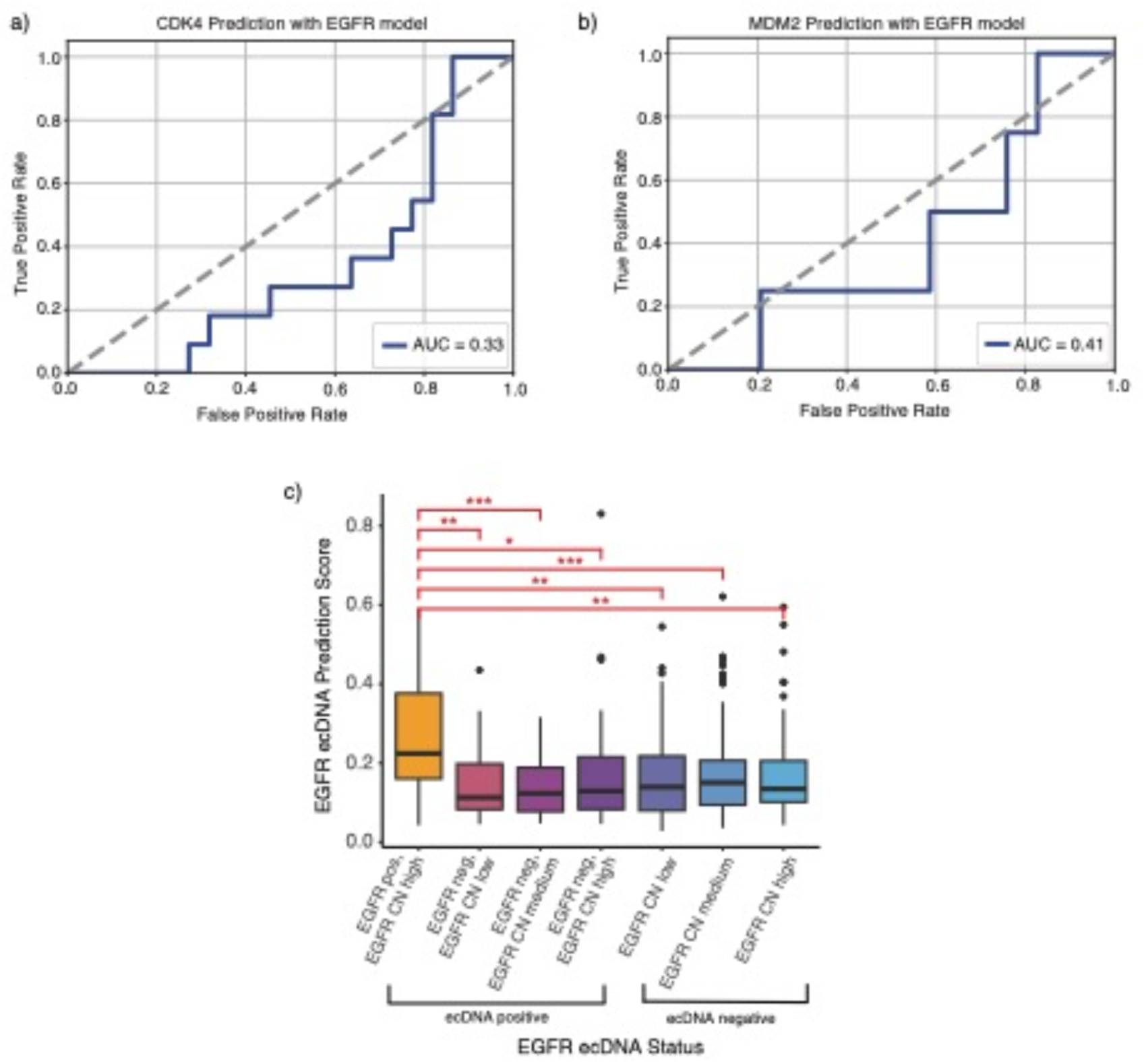
Evaluation of EGFR prediction model in CDK4, MDM2, and ecDNA negative samples. **a)** CDK4 status prediction ROC in LGG and GBM samples using the EGFR prediction model (Methods 5.2). **b)** MDM2 status prediction ROC in LGG and GBM samples using the EGFR prediction model. **c)** EGFR prediction probability in all ecDNA positive and ecDNA negative samples with known EGFR status. EGFR copy number (CN) obtained from TCGA is divided into low, medium, and high categories (low: normalized CN < 0, medium: 0 < normalized CN < 0.53, high: normalized CN > 0.53, Methods 1.8). EGFR pos: samples with ecDNA harboring EGFR oncogene; EGFR neg: samples with ecDNA that do not harbor the EGFR oncogene.

**Figure S8:**
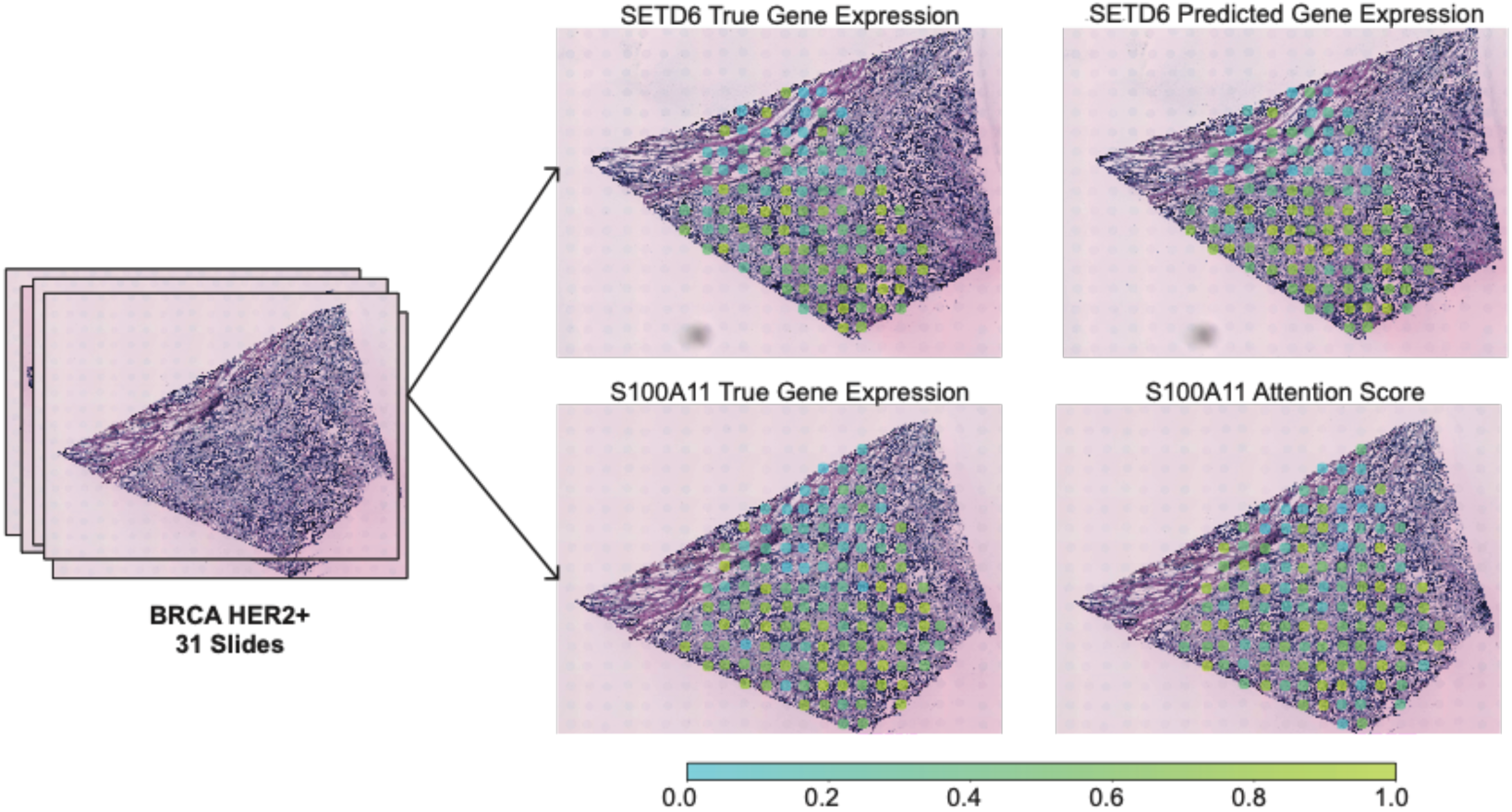
Visualizing predicted vs. true spatial expression variance of a few top ecPath feature. Visualization of rank normalized true gene expression, predicted gene expression, and attention scores (normalized between 0 and 1) for example BRCA patient in BRCA HER2+ slides. The top two slides and bottom two slides are from a single patients’ tissue that has had repeated H&E staining and microarray expression measures taken. Top example candidate gene (SETD6) with the highest concordance between true gene expression and predicted gene expression (Pearson R = 0.518, P =1.8e-08) and an example gene (S100A11) in the same patient with the highest concordance between true expression and attention score for gene expression prediction (Pearson R = 0.428, P = 6.5e-07) are shown.

**Figure S9.**
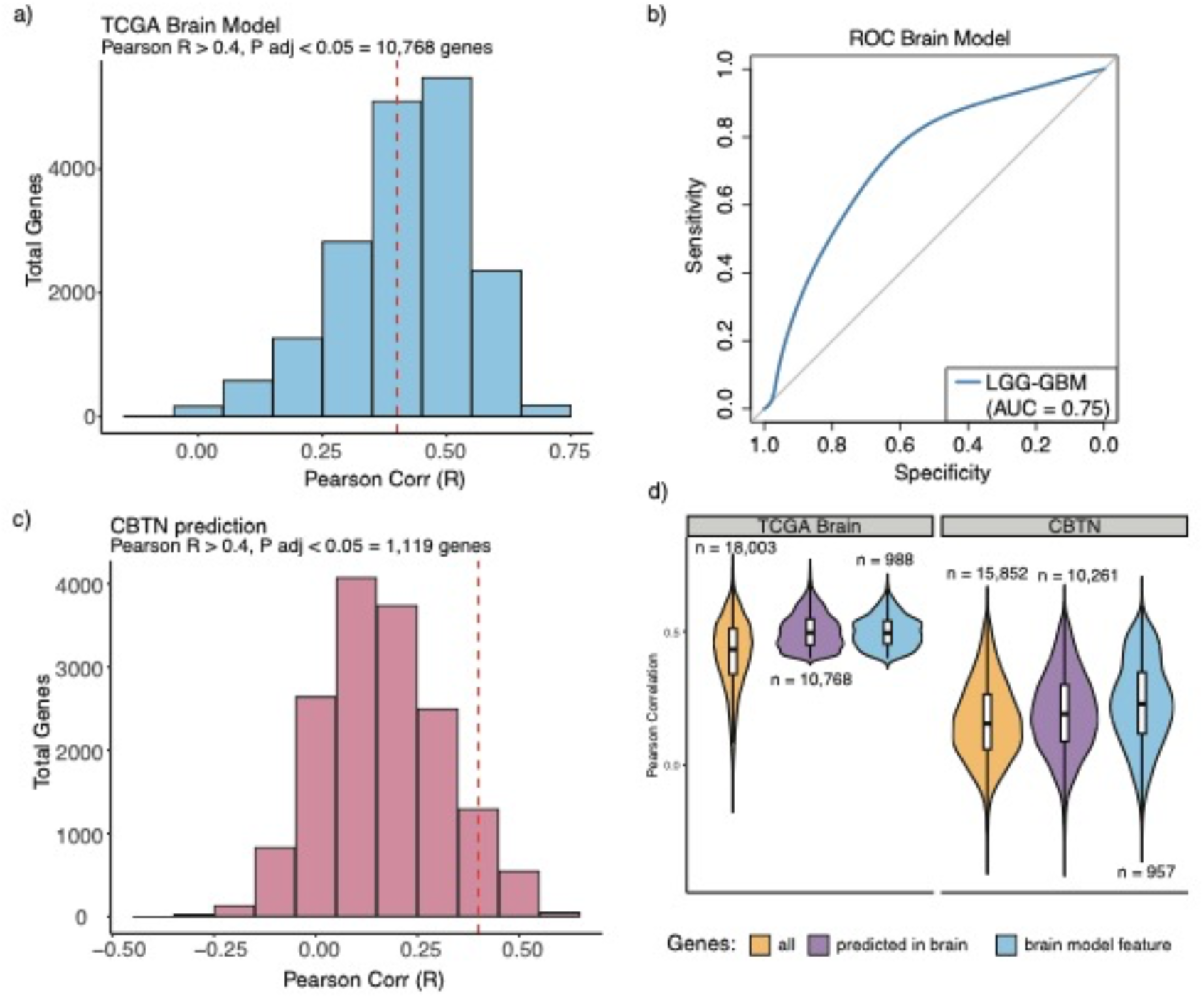
ecPath prediction performance of brain model on CBTN cohort. **a)** Pearson correlation distribution for true vs. predicted expression of all genes for LGG and GBM samples in the brain-specific logistic regression model. Dashed red line denotes a Pearson correlation of 0.4. **b)** ROC showing performance of the brain model on LGG and GBM samples from the logistic regression model. **c)** Distribution of Pearson correlation between measured CBTN gene expression and CBTN gene expression predictions from the TCGA brain model. Dashed red line denotes a Pearson correlation of 0.4. **d)** Pearson correlation of genes predicted by the TCGA brain-specific MLP model for LGG and GBM samples (left) or CBTN samples (right). Correlation values are shown for all genes (yellow), genes that can be significantly predicted by the brain model (purple) and those that are utilized in the subsequent logistic regression brain model to predict ecDNA status (blue).

**Table S1:** Total slides and patients in each of the 16 TCGA tumor types. analyzed in the ecPath MLP model.

| Tumor-type | slides | patients |
| --- | --- | --- |
| BRCA | 1,103 | 1,090 |
| KIRC | 529 | 524 |
| LGG | 524 | 506 |
| LUAD | 510 | 497 |
| HNSC | 494 | 492 |
| PRAD | 487 | 486 |
| LUSC | 485 | 485 |
| COAD | 466 | 454 |
| STAD | 365 | 365 |
| CESC | 306 | 304 |
| KIRP | 282 | 281 |
| PAAD | 178 | 177 |
| READ | 167 | 166 |
| ESCA | 151 | 150 |
| GBM | 150 | 147 |
| KICH | 65 | 65 |
| **Total** | **6,262** | **6,189** |

**Table S2:** Summary of ecPATH gene prediction from UNI across TCGA tumor types. Significance of gene expression predictions vs. TCGA measured expression were tested by Pearson correlation (R) and FDR corrected *P* value (Benjamin Hochberg).

| TUMOR-TYPE | TOTAL SAMPLES | TOTAL GENES REGRESSED | MEDIAN R | # GENES R > 0.4 | # GENES R > 0.75 | # SIGNIFICANT GENES (R > 0.4 & FDR 5%) |
| --- | --- | --- | --- | --- | --- | --- |
| LGG | 524 | 17,521 | 0.48 | 12,510 | 26 | 12,510 |
| KIRC | 529 | 17,496 | 0.43 | 10,255 | 108 | 10,255 |
| PRAD | 487 | 17,416 | 0.42 | 9,590 | 0 | 9,590 |
| KIRP | 282 | 17,174 | 0.41 | 8,922 | 42 | 8,922 |
| HNSC | 494 | 17,396 | 0.4 | 8,537 | 0 | 8,537 |
| STAD | 365 | 17,943 | 0.36 | 6,995 | 102 | 6,995 |
| LUAD | 510 | 17,435 | 0.37 | 6,930 | 39 | 6,930 |
| BRCA | 1103 | 17,400 | 0.36 | 6,814 | 33 | 6,814 |
| ESCA | 151 | 18,069 | 0.32 | 6,267 | 180 | 6,265 |
| PAAD | 178 | 17,570 | 0.34 | 6,159 | 0 | 6,159 |
| COAD | 466 | 16,894 | 0.35 | 5,792 | 2 | 5,792 |
| GBM | 150 | 17,546 | 0.32 | 5,355 | 2 | 5,355 |
| READ | 167 | 16,906 | 0.32 | 5,313 | 1 | 5,312 |
| LUSC | 485 | 17,641 | 0.33 | 5,167 | 0 | 5,167 |
| CESC | 306 | 17,272 | 0.31 | 4,709 | 3 | 4,709 |
| KICH | 65 | 16,963 | 0.26 | 3,902 | 21 | 3,875 |

**Table S3:** Samples for EGFR model. Total samples are those with known EGFR status, are ecDNA positive, and have gene expression predictions from ecPath. EGFR positive: ecDNA positive samples that harbor the EGFR oncogene, EGFR negative: ecDNA positive samples that do not harbor the EGFR oncogene.

| ***Tumor type*** | ***Total samples*** | ***EGFR positive*** | ***EGFR negative*** |
| --- | --- | --- | --- |
| ***LGG*** | 18 | 9 | 9 |
| ***GBM*** | 14 | 8 | 6 |
| ***HNSC*** | 35 | 4 | 31 |
| ***BRCA*** | 42 | 1 | 41 |
| ***ESCA*** | 11 | 1 | 10 |
| ***LUAD*** | 19 | 1 | 18 |
| ***Total*** | ***139*** | ***24*** | ***115*** |

**Table S4:** Pathologist interpretation of STK40 high attention regions for five example slides in the TCGA LGG cohort.

| Slide No. | Slide identifier | Tumor-type | Pathologist Interpretation of STK40 high attention regions |
| --- | --- | --- | --- |
| 1 | 44ac70e3 | LGG | higher density of nuclei, larger nuclei, more distinct cytoplasm |
| 2 | d41c157e | LGG | larger cells and larger nuclei |
| 3 | dd872474 | LGG | higher density of nuclei, larger and more pleomorphic nuclei, slightly more distinct cytoplasm |
| 4 | 158be9d2 | LGG | even distribution of tumor cells with less background artifact |
| 5 | 6ebd5bd4 | LGG | higher density of nuclei, larger and more pleomorphic nuclei, more distinct cytoplasm |

**Table S5:**
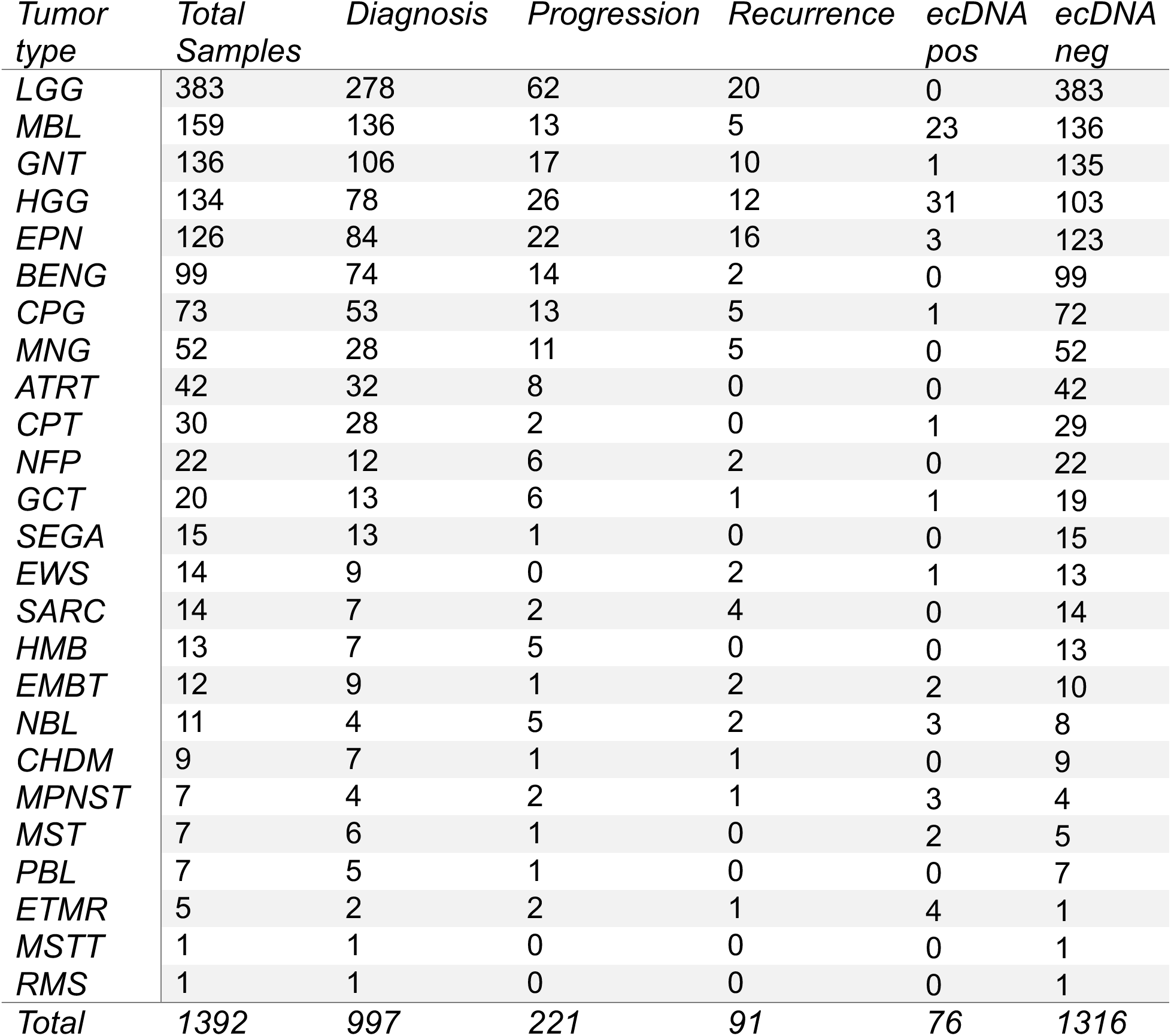
Child brain tumor network (CBTN) cohort overview. Total samples, samples in each diagnosis category, and ecDNA positive and negative samples are listed per tumor-type.

